# Discovery of a VHL molecular glue degrader of GEMIN3 by Picowell RNA-seq

**DOI:** 10.1101/2025.03.19.644003

**Authors:** Jonathan W. Bushman, Weixian Deng, Kusal T. G. Samarasinghe, Han-Yuan Liu, Shiqian Li, Amit Vaish, Alex Golkar, Shu-Ching Ou, Jinwoo Ahn, Rajesh K. Harijan, Han Han, Tomas Duffy, Elliot Imler, Judy Wu, Renee Rawson, Lisa Liu, John Maina Ndungu, Michael D. Hocker, Animesh Roy, Amber D. Senese, Chun-Chieh Lin, Duc Tran, Alex Campos, Gabrielle Blanco, Julia I. Toth, Anastasia Velentza, Andrew Burritt, Gregory S. Parker, Peggy A. Thompson, Simon Bailey, Willem den Besten, Stephanie Voss, Bo Zhou, Kate S. Ashton, Olivier Bedel, Jaeki Min, Patrick Ryan Potts

## Abstract

Targeted protein degradation (TPD) is an emerging therapeutic modality in which small molecules are used to recruit targets to the natural protein degradation machinery of the cell. Molecular glue degraders (MGD) are monovalent small molecules that accomplish this by redirecting E3 ubiquitin ligases to target proteins, offering the potential to degrade previously unliganded and “undruggable” proteins in cancer, neurodegenerative, and other diseases. While attractive due to their drug-like properties, MGDs are exceptionally hard to discover and have largely been identified serendipitously. The Von Hippel-Lindau (VHL) E3 ligase is the second most widely used effector for TPD, though current VHL-based degraders are primarily large heterobifunctional PROTACs (proteolysis-targeting chimeras) designed using target-based ligands. Here, we have instead pursued target-agnostic discovery of VHL MGDs leveraging proprietary ultra-miniaturized microfluidics devices (Picowells) to facilitate unbiased RNA-seq screening of a biased E3-focused library. This resulted in dGEM3, a novel VHL molecular glue that targets the survival of motor neuron (SMN) complex member GEMIN3 for degradation. Through a combination of cellular, biochemical, and biophysical assays, we have characterized the GEMIN3 degron within its helicase ATP-binding domain, and how the kinetics of ternary complex formation impact degradation. These findings provide insights on the re-programmability of VHL for novel targets using drug-like molecular glues.

## Introduction

Targeted protein degradation (TPD) refers to the use of proximity-inducing molecules that redirect target proteins to the natural degradation machinery of the cell. The earliest examples of TPD used heterobifunctional molecules called PROTACs (proteolysis-targeting chimeras) to bring a protein of interest to an E3 ubiquitin ligase, which can ubiquitinate and send the target to the proteasome for degradation^1-4^. These degrader molecules are rationally designed by chemically linking a ligand for a target of interest to an E3 ligase ligand. As one of the first E3 ubiquitin ligases with a structurally validated small molecule ligand, the Von Hippel-Lindau (VHL) E3 ligase saw early widespread adoption in the development of PROTACs as chemical biology tools^5^. More recently, VHL-based molecules have entered clinical investigation for therapeutically relevant targets^6-10^. However, the ongoing development of PROTACs faces challenges from the large size and poor drug-like properties of these molecules^11^.

Molecular glue degraders (MGDs) provide an alternative chemical approach to TPD, instead using monovalent small molecules that bind one protein and enhance existing protein-protein interactions (PPIs) or induce neomorphic PPIs, redirecting a target for degradation. Molecular glues have several inherent advantages over heterobifunctional molecules, including avoiding the “hook effect” caused by saturation of bivalent binding at high concentrations, as well as improved drug-like properties related to their lower molecular weight. However, the discovery and rational design of MGDs has proven especially challenging and is still limited to a small fraction of the approximately 600 E3 ligases encoded in the human proteome^7,12^. Discovery of molecular glues has predominantly occurred after careful investigation of inhibitor ligands with known phenotypic effects, including immunomodulatory (IMiD) drugs^13-19^, anticancer aryl sulfonamides^20-24^, cytotoxic pan-CDK kinase inhibitors^25-29^, and a HIF-1α-stabilizing VHL inhibitor^30,31^. While the serendipitously discovered MGDs from these examples are all E3 ligase ligands (for CRBN, DCAF15, DDB1, and VHL respectively), emerging evidence suggests that monovalent degraders derived from target-based inhibitors can also be used for TPD. Recently discovered molecules based on inhibitors of the BCL6 BTB domain, BRD4 bromodomain, and various kinases have revealed molecular glues that recruit SIAH1, DCAF16, and DCAF11^32-41^.

Despite these revelations, there remain significant challenges in identifying additional E3 ligase ligands for TPD, especially molecules that will expand the target space for protein degradation by molecular glues. Although CRBN and VHL have both proven effective for PROTAC development, CRBN has so far emerged as the most prominent E3 ligase for MGD development, with VHL molecular glues limited to a single example targeting CDO1^30^. To address these challenges, we took inspiration from the success of developing CRBN MGDs by expanding on the IMiD scaffold, as well as the success of recent target-agnostic rational molecular glue screens^11,27,42-47^. We leveraged extensive existing chemistry for VHL inhibitors to identify new molecular glues and targets for the second most widely used E3 ligase for TPD. In this work, we report the discovery of the VHL molecular glue dGEM3 using ultra-high throughput Picowell RNA-seq screening, a unique molecular glue screening approach combining a biased E3 ligase-focused, DNA-encoded chemical library (DEL) with an unbiased RNA-seq readout. We reveal GEMIN3 as a neosubstrate of VHL that is targeted for degradation by its helicase ATP-binding domain, with biophysical studies of the molecular glue confirming the importance of stable ternary complex formation for TPD. We also identified the critical interaction domains of GEMIN3 through studies involving rational truncation, mutation, and domain swapping using cellular ternary complex formation.

## Results

### Discovery of dGEM3 from a cellular VHL-focused DEL screen

Previous efforts to discover MGDs have seen success with multiple approaches, including the conversion of target-based inhibitors into degraders by chemical derivatization, as well as phenotypic screens to identify hit molecules regardless of the degraded target^27,29,33,47^. We hypothesized that constructing small molecule libraries based on ligands for the VHL E3 ligase and screening in a target-agnostic fashion would result in the identification of novel VHL MGDs. This library design incorporated structural features which are known to be important for binding to VHL based upon reported ligands^5^. Putative VHL binding motifs were then elaborated at multiple exit vectors to maximize the diversity of the ligand-bound VHL surface presented for protein-protein contacts with potential targets^48,49^. The bead-based library design also made use of a photocleavable linker to allow cellular treatment, with compounds released from the beads unencumbered by a DNA barcode and able to enter cells.

With VHL-focused libraries in hand, we aimed to identify active molecules in a target-agnostic manner using Picowell RNA-seq. Given that degradation of cellular proteins should lead to altered signaling and cellular states that can be inferred by differential gene expression profiles, high-throughput Picowell RNA-seq screens encompassing approximately 26,000 molecules were conducted to identify hits that induced differential gene expression after 24 hour treatment (Fig. 1a). From a sub-library of 4,327 molecules, 28 initial hits were determined to significantly alter the cellular gene expression profile and were resynthesized off-bead for follow-up validation by cellular VHL binding assay and bulk RNA-seq (Fig. 1b). Consistent with the library design, most of the hits demonstrated binding to VHL, with 18 compounds competitively displacing a VHL tracer with an IC_50_ below 10 µM (Fig. 1c). Bulk RNA-seq revealed a wide range of cellular activities, with several compounds inducing no differentially expressed genes (DEGs) and four compounds perturbing greater than 1,000 DEGs (Fig. 1c).

**Fig. 1.**
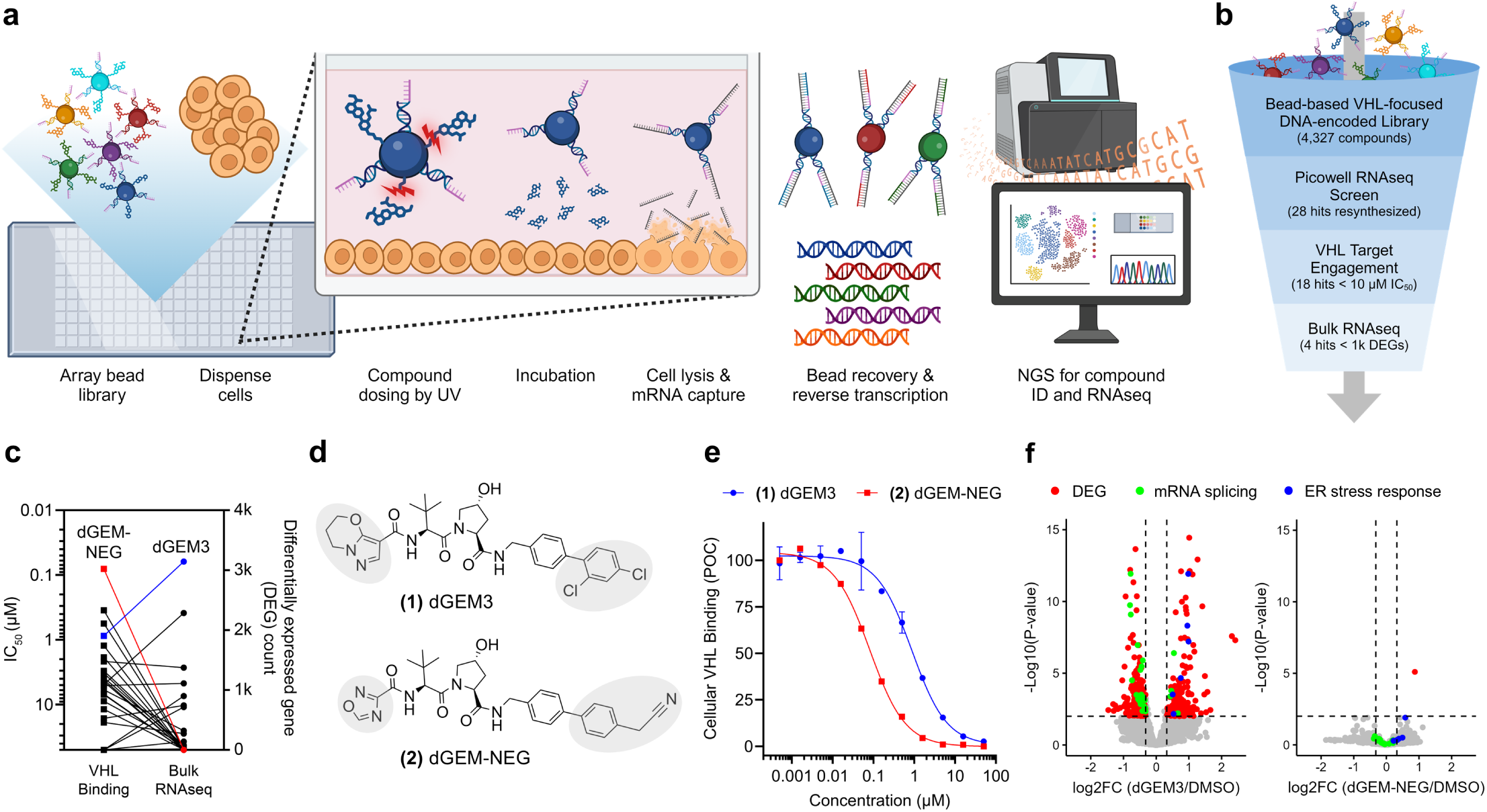
DEL-based Picowell RNA-seq screen identifies dGEM3 as a VHL ligand that perturbs the transcriptome. **a,** Picowell screening platform for RNA-seq screening of bead-based compound libraries **b**, Screening and validation workflow to identify VHL molecular glues from a bead-based, VHL-focused DEL. DEL, DNA-encoded library; DEG, differentially expressed gene; IC_50_, half-maximal inhibitory concentration. **c**, Validation of primary screen by VHL NanoBRET target engagement assay and 24 h bulk RNA-seq **d**, Chemical structures of dGEM3 and dGEM-NEG. Differences from common VHL ligands (VH032 and VH298) are shaded grey. **e,** VHL NanoBRET target engagement assay measuring displacement of VHL tracer by competition with increasing concentrations of hit compound for 2 h in HEK293T cells transfected with NanoLuc-VHL. **f**, Bulk RNA-seq validation of dGEM3 and dGEM-NEG following 6 h treatment at 10 μM in HEK293T cells. Dashed lines indicate significance and enrichment cutoffs of p-value < 0.01 and FC > 1.25. Genes corresponding to top enriched up- and down-regulated gene ontology terms shaded in blue and green, respectively.

Two compounds were selected for in-depth characterization—dGEM3 as the most active compound by RNA-seq and dGEM-NEG as the highest-affinity VHL binder from the resynthesized hits (Fig. 1d). Both dGEM3 and dGEM-NEG preserve the central hydroxyproline motif from VHL ligands based on the endogenous VHL substrate HIF-1α^5^. However, dGEM3 is characterized by a unique right-hand-side (RHS) dichlorobenzene and left-hand-side (LHS) pyrazolo-oxazine heterocycle. dGEM-NEG contains a related RHS biphenyl group, replacing the chlorine atoms with a single cyanomethyl group instead and substituting the LHS for an oxadiazole. Both compounds demonstrated sub-micromolar binding to VHL, with IC_50_ values of 870 nM and 80 nM for dGEM3 and dGEM-NEG, respectively (Fig. 1e). Curiously, the two compounds demonstrated dramatically different cellular activity. Treatment with 10 µM dGEM3 induced broad changes to the transcriptome after 6 hours, including both up-regulation of genes involved in the ER stress response and down-regulation of genes involved in mRNA splicing (Fig. 1f). In contrast, dGEM-NEG caused no significant changes to the transcriptome aside from upregulation of the ER stress response gene TRIB3.

### GEMIN3 is an SMN complex subunit targeted for degradation by dGEM3

To investigate the source of the transcriptional signature caused by dGEM3, we utilized a proximity labeling proteomics approach, TurboID^50,51^, to assess whether dGEM3 could induce proximity between VHL and any potential cellular targets regulating gene expression (Fig. 2a). Indeed, TurboID proteomics indicated that dGEM3 recruits subunits of the survival of motor neuron (SMN) complex to VHL, including SMN2, GEMIN3, GEMIN4, GEMIN5, GEMIN6, and GEMIN8 (Fig. 2b, S1a). The SMN complex is responsible for biogenesis of spliceosomal small nuclear ribonucleoproteins (snRNPs) that play a key role in mRNA splicing and are essential for cell survival, highlighted by the central role of this complex in the neurodegenerative diseases amyotrophic lateral sclerosis (ALS) and spinal muscular atrophy (SMA)^52^.

**Fig. 2.**
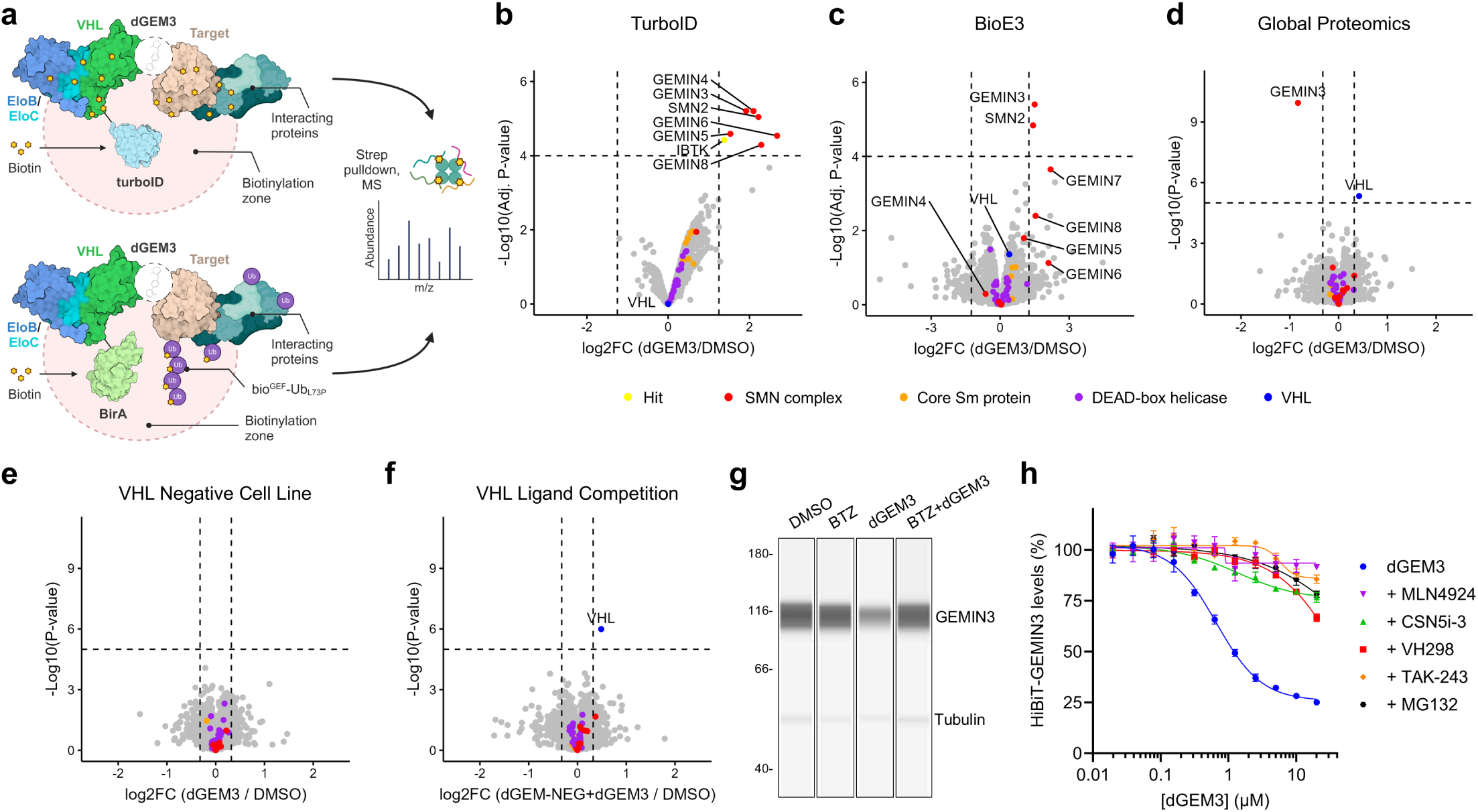
GEMIN3 is the SMN complex subunit targeted to the ubiquitin proteasome system for degradation by dGEM3. **a**, Schematic of enrichment-based proteomics approaches, TurboID (top) and BioE3 (bottom). The TurboID cell line expresses VHL fused to the TurboID biotin ligase that biotinylates proteins in proximity to VHL, which may be enriched by pull-down and identified by mass spectrometry. BioE3 uses HEK293 VHL KO cells expressing VHL-BirA and bio^GEF^-Ub_L73P_ for site-specific labeling of ubiquitinated proteins proximal to VHL following biotin pulse. **b**, TurboID-VHL proximity labeling assay in K-562 cells treated for 6 h with 10 μM dGEM3 (n=3) or 0.1% DMSO (n=3) and biotin, following 1 h pre-treatment with proteasome inhibitor MG132. Dashed lines indicate significance and enrichment cutoffs of adjusted p-value < 0.0001 and LFC > 1.25. Interaction partners (SMN complex, red; Core Sm proteins, orange) and family members (DEAD-box helicases, purple) of enriched hit proteins are indicated by color. LFC, Log_2_ Fold Change. **c**, BioE3 VHL proximity ubiquitin labeling assay following co-treatment with 10 μM dGEM3 (n=3) or 0.1% DMSO (n=3) and 2 μM BTZ for 2 h. Significance, enrichment cutoffs, and legend the same as Fig. 2b. BTZ, bortezomib. **d**, Global proteomics following treatment with 10 μM dGEM3 (n=4) or 0.1% DMSO (n=4) for 6 h in HEK293T cell lines. Dashed lines indicate significance and enrichment cutoffs of p < 0.00001 and FC > 1.25. Legend same as Fig. 2b. See Fig. S2b for additional cell lines and 24 h time points. **e**, Global proteomics of dGEM3 treatment in VHL-deficient 786-O cancer cell line. Replicates, cutoffs, and colors are identical to Fig. 2d. See Fig. S2c for 24 h time point. **f**, Global proteomics of dGEM3 treatment following pre-treatment with competitor dGEM-NEG. Replicates, cutoffs, and colors are identical to Fig. 2d. See Fig. S2d for 24 h time point. **g**, Simple Western analysis of GEMIN3 protein levels following treatment with dGEM3. BTZ alone and BTZ co-treatment confirm proteasome dependence. **h**, Mechanism-of-action evaluation of GEMIN3 degradation by dGEM3. HiBiT-GEMIN3 cells were treated in dose-response with dGEM3 for 15 h, following 1 h pre-treatment with the indicated inhibitors of ubiquitin-proteasome system components. Pre-treatments included inhibitors of ubiquitin-activating enzyme E1 (1 μM TAK-243), CRL neddylation cycle (1 μM MLN4924, 1 μM CSN5i-3), proteasome (10 μM MG132), and VHL (50 μM VH298). Mean ± s.e.m.; n = 4.

As E3 ligases are known to ubiquitinate target proteins in a proximity-dependent manner, we hypothesized that SMN complex recruitment to VHL would lead to ubiquitination and degradation of its protein subunits by the ubiquitin proteasome system (UPS). To establish which proteins are directly ubiquitinated by VHL following dGEM3 treatment, we utilized BioE3, a proximity labeling proteomics approach for site-specific biotinylation and enrichment of ubiquitinated substrates of a tagged E3 ligases^53,54^ (Fig. 2a). BioE3 proteomics revealed that the GEMIN3 and SMN2 subunits of the SMN complex are ubiquitinated by VHL following dGEM3 treatment, while other subunits of the SMN complex were quantified but not significantly affected (Fig. 2c).

To assess whether dGEM3 treatment leads to degradation of SMN2 and GEMIN3 or any other potential targets, we performed unbiased global proteomics in multiple cancer cell lines. Consistent with the effects of previously reported VHL inhibitors^55^ as well as dGEM-NEG (Fig. S2a), VHL itself was slightly up-regulated following dGEM3 treatment (Fig 2d). Remarkably, out of 9,877 quantified proteins across five cell lines (HEK293T, MV-4-11, MCF-7, JVM-3, and SH-SY5Y), only a single protein, GEMIN3, was significantly degraded after 6 hour dGEM3 treatment (Fig. 2d, S1b). GEMIN3, also known as DDX20 or DP103, is the DEAD-box helicase subunit of the SMN complex that unwinds small nuclear RNAs (snRNAs) during maturation of spliceosomal snRNPs^56,57^. Notably, SMN2 protein abundance was not affected in any of the tested cell lines. To confirm whether GEMIN3 degradation is dependent on VHL, dGEM3 treatment was assessed in a VHL-deficient cell line. Global proteomics revealed that dGEM3 does not impact GEMIN3 protein levels in 786-O cells, confirming that GEMIN3 degradation is indeed dependent on VHL (Fig. 2e, S1c). Degradation of GEMIN3 could also be prevented by ligand competition with dGEM-NEG, indicating that dGEM3 recruits GEMIN3 for degradation through the expected VHL HIF-1α binding pocket (Fig. 2f, S1d, S2b). Supporting the mass spectrometry analysis, the 50% GEMIN3 degradation seen by global proteomics in HEK293T cells was also confirmed by simple western analysis (Fig. 2g).

To further assess the depth and potency of target degradation by dGEM3, a cellular HiBiT GEMIN3 degradation assay was established by knocking in a HiBiT tag for protein quantification to the endogenous GEMIN3 gene in HEK293T cells. Following GEMIN3 protein levels in response to dGEM3 treatment over time revealed that GEMIN3 was degraded with a maximal degradation (D_max_) of 50% at 6 hours and 75% at 18 hours (Fig. S1e). These D_max_ values were consistent with the protein abundance changes observed at both time points by global proteomics (Fig. S1b). dGEM3 also demonstrated dose-dependent degradation of GEMIN3 with a half-maximal degradation concentration (DC_50_) of 680 nM (Fig. 2h). Importantly, GEMIN3 degradation was abolished by inhibitors of multiple components of the UPS, including proteasome inhibitor (MG132), E1 ubiquitin-activating enzyme inhibitor (MLN7243), and VHL inhibitor (VH298) (Fig. 2h). As VHL is the substrate receptor of a CUL2 RING E3 ubiquitin ligase (CRL2) complex, which requires dynamic neddylation for E3 ligase activity, we also tested whether inhibitors of the CRL neddylation cycle (MLN4924, CSN5i-3) would abolish GEMIN3 degradation, which they did (Fig. 2h). Taken together, these results establish that dGEM3 selectively targets GEMIN3 in a VHL-dependent manner for targeted protein degradation through the UPS. Having confirmed GEMIN3 as the sole protein targeted for degradation, we designate this molecular glue degrader as dGEM3, or **d**egrader of **GEM**IN**3**.

### dGEM3 is a molecular glue between VHL and the GEMIN3 helicase ATP-binding domain

Having established that dGEM3 redirects VHL to degrade GEMIN3, we sought to understand the structural basis of substrate recognition by VHL and the proteome-wide selectivity of dGEM3. We anticipated that dGEM3 acts as a molecular glue between VHL and a structural degron unique to GEMIN3 among the DEAD-box helicase family. To facilitate identification of a minimal GEMIN3 construct for further study, a cellular NanoBiT ternary complex assay was established using a split luciferase complementation system for detecting chemically induced proximity of VHL and GEMIN3 (Fig 3a). The tag orientation of VHL and GEMIN3 was optimized to rule out experimental artifacts from tagging, with three of the four tested configurations showing dGEM3-dependent ternary complex formation (TCF) between VHL and full-length GEMIN3 (Fig. 3b). To confirm whether this TCF was dependent on dGEM3 binding in the expected HIF-1α binding pocket of VHL, the dGEM3 dose-response was repeated with competition using 10 µM of the VHL inhibitor VH298^31^. This resulted in an approximately 10-fold increase in EC_50_ (0.58 µM vs 5.72 µM) of dGEM3 with full-length GEMIN3, confirming that VH298 and dGEM3 act at the same binding pocket (Fig. 3c). VHL therefore is likely interacting with GEMIN3 using an interface proximal to that with which it recognizes its endogenous substrate HIF-1α.

**Fig. 3.**
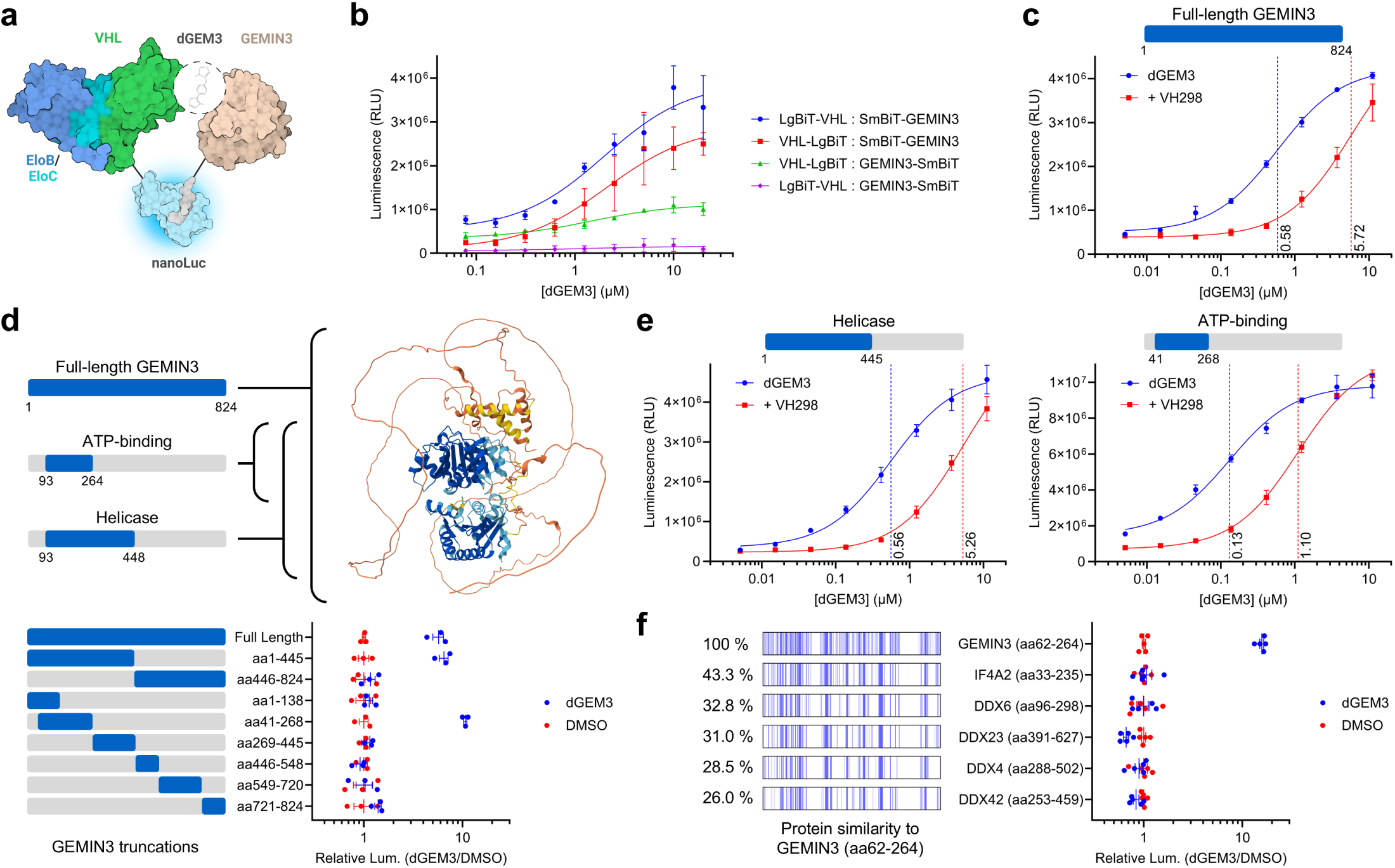
dGEM3 is a selective molecular glue between VHL and the GEMIN3 ATP-binding domain. **a**, Schematic of cellular VHL ternary complex formation assay. Split-luciferase complementation is achieved by chemically induced proximity of LgBiT-tagged VHL and SmBiT-tagged target protein. **b**, Optimization of tag orientation for VHL ternary complex formation assay using VHL and full-length GEMIN3. HEK293T cells expressing LgBiT-tagged VHL and SmBiT-tagged GEMIN3 were treated with dGEM3 in dose-response for 2 h after 0.5 h treatment with 1 μM BTZ. Mean ± s.e.m.; n = 3. **c**, Full-length GEMIN3 VHL ternary complex assay with dGEM3 dose-response and VHL ligand competition. HEK293T cells expressing VHL-LgBiT and SmBiT-GEMIN3 were treated with dGEM3 for 2 h following pre-treatment with 1 μM BTZ and 10 μM VH298 or DMSO. Dashed lines indicate the EC_50_ for respective dose-response curves by color. Mean ± s.e.m.; n = 3. **d**, GEMIN3 truncation analysis by VHL ternary complex assay. HEK293T cells expressing VHL-LgBiT and SmBiT-GEMIN3 truncation constructs were pre-treated with BTZ and subjected to treatment with 10 μM dGEM3 or DMSO for 2 h. Active constructs correspond roughly to the full-length, helicase domain, and ATP-binding domain as indicated by the AlphaFold structure prediction. Luminescence ratio was calculated relative to DMSO control. Mean ± s.e.m.; n = 3. **e**, GEMIN3 helicase and ATP-binding domain VHL ternary complex assay with dGEM3 dose-response and VHL ligand competition. HEK293T cells expressing VHL-LgBiT and SmBiT-GEMIN3_aa1-445_ or SmBiT-GEMIN3_aa41-268_ were treated with dGEM3 for 2 h following pre-treatment with 1 μM BTZ and 10 μM VH298 or DMSO. Dashed lines indicate the EC_50_ for respective dose-response curves by color. Mean ± s.e.m.; n = 3. **f**, Selectivity for DEAD-box helicases by VHL ternary complex assay. Alignment of constructs is based on the helicase ATP-binding domain and shaded by percent protein sequence identity to GEMIN3. Cells pre-treated with BTZ were subjected to 2 h 10 μM dGEM3 or DMSO treatment and luminescence was calculated per-construct relative to DMSO controls. Mean ± s.e.m.; n = 5. See Fig. S3 for full alignment.

To establish which part of GEMIN3 is recognized by VHL, rational truncation constructs of GEMIN3 were designed based on the AlphaFold structure prediction and prior studies^58-61^. GEMIN3 consists of an N-terminal DEAD-box helicase domain, followed by an unstructured C-terminus predicted to mediate protein-protein interactions^56^. Three overlapping constructs (GEMIN3_aa1-824_, GEMIN3_aa1-445_, and GEMIN3_aa41-268_) demonstrated clear dGEM3-dependent TCF with VHL, corresponding roughly to full-length GEMIN3, the helicase domain, and the helicase ATP-binding domain (Fig. 3d). These minimal constructs were sufficient for TCF in dose-response, with respective EC_50_ values of 560 nM for the helicase and 130 nM for the ATP-binding domain. Like the full-length construct, VH298 competition confirmed that TCF between VHL and the helicase or ATP-binding domains of GEMIN3 was dependent on the HIF-1α binding pocket of VHL (Fig. 3e). These results suggest that the ATP-binding domain of GEMIN3 alone is sufficient for cellular TCF and the degron responsible for GEMIN3 degradation lies in this domain.

With a minimal construct identified, we sought additional potential targets of dGEM3 by using the Foldseek webserver to search for proteins that share structural similarity to crystal structures of the GEMIN3 ATP binding domain (GEMIN3_aa41-268_) (PDB: 2OXC, 3B7G)^62,63^. The majority of potential candidates belonged to the 60-member human DEAD-box helicase family, with sequence identity ranging from 20 to 40 percent. A subset of candidate ATP-binding domains was tested for TCF with VHL, with GEMIN3 remaining the only helicase ATP-binding domain recruited by dGEM3 (Fig. 3f). Taken together with the TurboID and global proteomics experiments (Fig. 2b, 2d), these results confirm the exquisite selectivity of the molecular glue for GEMIN3 in both ternary complex formation and degradation.

### Biochemical reconstitution of the VHL:dGEM3:GEMIN3 ternary complex

While the prior findings in cellular assays strongly suggest that dGEM3 is a molecular glue between VHL and the GEMIN3 helicase ATP-binding domain, full biochemical reconstitution of the ternary complex was necessary to fully characterize the molecular glue activity. Using purified VHL/EloB/EloC (VBC) complex, GEMIN3_aa41-268_, and BRD4_BD2_ (as a control), we established an *in vitro* homogenous time-resolved fluorescence (HTRF) assay capable of detecting molecular glue or PROTAC-induced TCF (Fig. 4a). Consistent with cellular data, dGEM3 induced a ternary complex between purified VHL and the GEMIN3 ATP-binding domain with an EC_50_ of 450 nM (Fig. 4b). The positive control MZ1 demonstrated TCF between VHL and BRD4 with a prominent hook effect characteristic of heterobifunctional PROTACs. In contrast, dGEM3 showed no evidence of hook effect, consistent with a molecular glue mechanism of action. As expected, the VHL inhibitors VH298 and VH032 were both unable to recruit GEMIN3 to VHL. Confirming the specificity of the dGEM3-induced interaction between VHL and GEMIN3, the ternary complex was also reversible by counter-titration of VH298 (Fig. 4c).

**Fig. 4.**
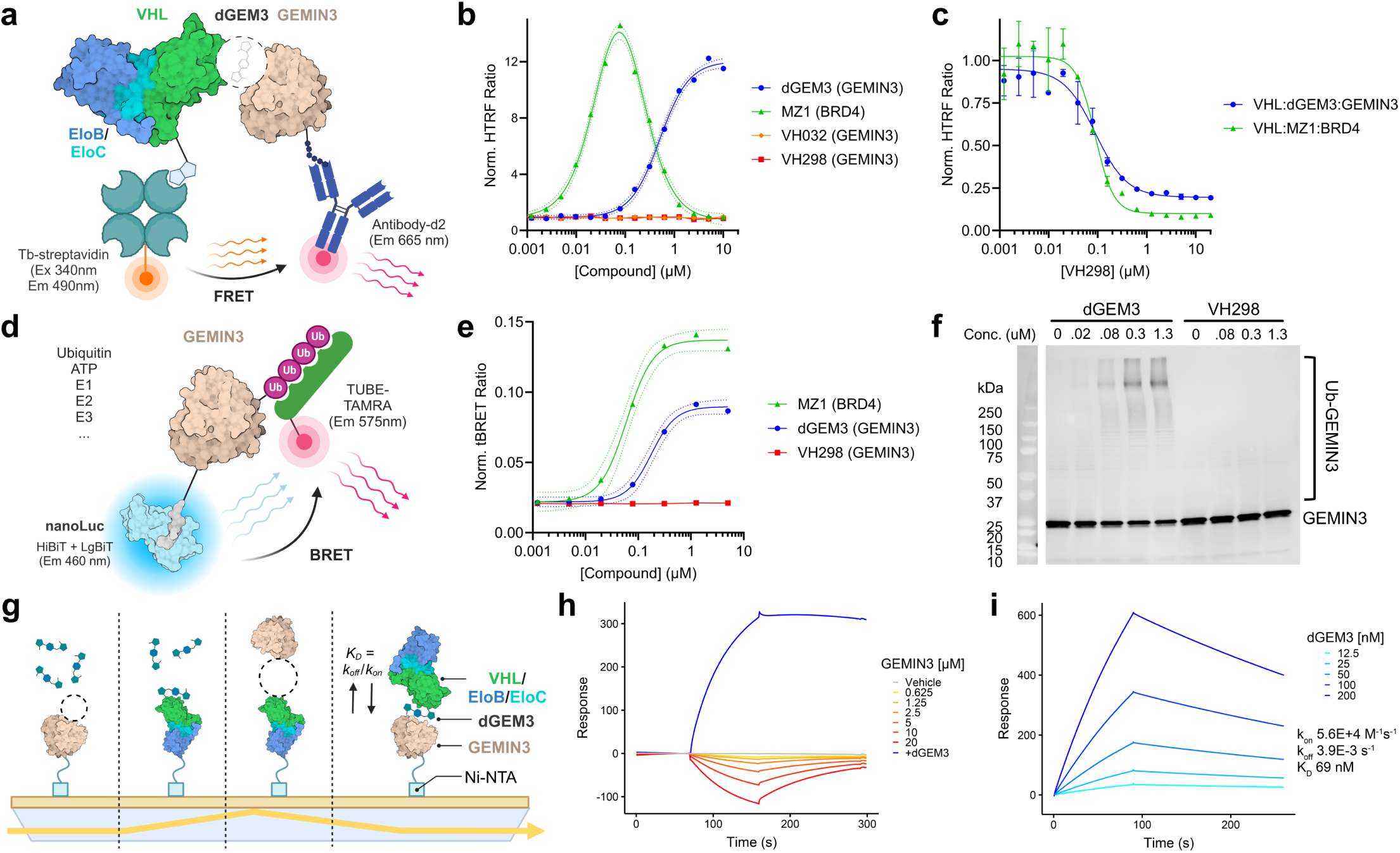
Biochemical reconstitution of the VHL:dGEM3:GEMIN3 ternary complex and ubiquitination. **a**, Schematic of HTRF *in vitro* VHL ternary complex assay. Biotinylated AviTag-VHL/EloB/EloC complex (VBC) was incubated with His-tagged GEMIN3_aa41-268_ or BRD4_BD2_ control, and Streptavidin-Tb with MAb Anti-6His-d2 for detection of ternary complex formation by HTRF. **b**, *In vitro* VHL ternary complex formation of dGEM3 and VHL-based PROTAC and inhibitors. VHL was incubated with indicated target and compound for 1 h as in Fig. 4a. Solid lines indicate regression curve fit to 14-point 2-fold dose-response, and dashed lines indicate 95% confidence bands for nonlinear fit. **c**, *In vitro* VHL ternary complex competition assay. TCF was induced by co-incubation of 10 nM VBC with 1 μM dGEM3 and 40 nM GEMIN3_aa41-268_, or 50 nM MZ1 and 40 nM BRD4_BD2_. VH298 was added in dose response before the 1 h incubation to competitively disrupt TCF. Mean ± s.e.m.; n = 2. **d**, Schematic of tBRET *in vitro* ubiquitination assay with purified Ub, UBA1, UBE2L3, ARIH1, UBE2R2, N8-CRL2-EloB-EloC-VHL, and HiBiT-GEMIN3_aa41-268_ or BRD4_BD2_ control. Detection is achieved by BRET between the complemented HiBiT-target split-luciferase and TAMRA-linked tandem ubiquitin-binding entities (TUBE). **e**, tBRET *in vitro* ubiquitination assay for dGEM3-dependent ubiquitination of GEMIN3_aa41-268_ and MZ1-dependent ubiquitination of BRD4_BD2_. Compounds were incubated with their respective targets in the ubiquitination reaction for 0.5 h before detection by tBRET or immunoblot. Solid lines indicate regression curve fit to 8-point 4-fold dose-response, and dashed lines indicate 95% confidence bands for nonlinear fit. **f**, HiBiT immunoblot of *in vitro* ubiquitination reaction for GEMIN3_aa41-268_. **g**, Schematic of SPR assays for dGEM3 molecular glue characterization. His-tagged VBC or GEMIN3_aa41-268_ was immobilized on a Ni-NTA chip and used to evaluate 1) GEMIN3:compound binding, 2) VHL:compound binding, 3) VHL:GEMIN3 binding without compound, and 4) VHL:dGEM3:GEMIN3 ternary complex formation. **h**, Absence of interaction between VHL and GEMIN3 by SPR. GEMIN3_aa41-268_ was titrated against immobilized VBC up to 20 μM, with no specific interaction observed. Negative response indicates non-specific binding to the reference cell. 400 nM dGEM3 was included as positive control. **i**, VHL:dGEM3:GEMIN3 ternary complex formation by SPR. The VBC:dGEM3 complex was pre-formed and serially diluted before analysis on a low-density GEMIN3-functionalized surface. Data was fit to a 1:1 Langmuir interaction model and equilibrium ternary complex binding affinity was calculated using a steady-state affinity model. K_on_, association rate constant; K_off_, dissociation rate constant; K_D_, equilibrium dissociation constant.

Based on the observed proteasome-dependent degradation of GEMIN3 (Fig. 2e), we reasoned that it would be possible to reconstitute ubiquitination of GEMIN3 by VHL. We established an *in vitro* bioluminescence resonance energy transfer (BRET) assay to quantitate GEMIN3 ubiquitination utilizing a tandem ubiquitin-binding entity (TUBE)-TAMRA reagent to detect substrate polyubiquitin chains. Initial attempts to demonstrate GEMIN3_aa41-268_ ubiquitination using E1 (UBA1), E2 (UBE2R2), and active neddylated E3 CRL2^VHL^ were unsuccessful. However, supplementing VHL with the priming E3 ligase ARIH1 and its cognate E2 UBE2L3^64,65^ resulted in robust dGEM3-dependent polyubiquitination of GEMIN3 (Fig. 4e). Polyubiquitination of GEMIN3 was also detectable by western blot (Fig. 4f). VH298 was included as a negative control to confirm GEMIN3 ubiquitination is specific to dGEM3 and not a byproduct of promiscuous enzymatic activity in the *in vitro* reaction.

Molecular glue degraders differ from heterobifunctional PROTACs in that they only show appreciable binding to one member of the ternary complex. It is also widely speculated that molecular glues may exploit weak endogenous interactions between E3 and substrate to form productive ternary complexes. We used surface plasmon resonance (SPR) for biophysical characterization of the VHL:dGEM3:GEMIN3_aa41-268_ ternary complex, defining key features of the molecular glue interaction (Fig. 4g). First, we used GEMIN3- or VBC-functionalized surfaces to probe binary binding of dGEM3. dGEM3 displayed monovalent binding to VHL (K_D_ = 217 nM) but not GEMIN3. Second, we investigated potential binding between immobilized VHL and free GEMIN3 up to 20 µM in the absence of compound. No weak, basal interaction could be detected between GEMIN3 and VHL, suggesting that the two proteins do not interact at relevant cellular concentrations (Fig. 4h). Lastly, we measured the association and dissociation kinetics of the VHL:dGEM3:GEMIN3_aa41-268_ ternary complex, calculating a dissociation constant (K_D_) of 69 nM (Fig. 4i). Compared to previous VHL PROTACs, dGEM3 demonstrates much slower kinetics, with both slower association (k_on_ = 5.6E+4) and dissociation (k_off_ = 5.6E+4 M^-1^s^-1^) rates^66^. The totality of the biochemical and biophysical data supports the notion that dGEM3 is a *bona fide* VHL molecular glue for the GEMIN3 helicase ATP-binding domain.

### Mapping the GEMIN3 degron based on species selectivity of dGEM3

We sought to leverage species selectivity among GEMIN3 orthologs to narrow down a region of the helicase ATP-binding domain responsible for ternary complex formation with VHL. While the ATP-binding domains of human DEAD-box helicases possess less than 50% similarity to GEMIN3 and are not recruited by dGEM3 (Fig. 3f), GEMIN3 orthologs possess much higher homology (Fig. S3). Orthologs of the ATP-binding domain from *Mus musculus* and *Xenopus tropicalis* GEMIN3 were tested for TCF by cellular NanoBiT assay, confirming dGEM3-dependent TCF in both mouse and frog (Fig. 5a). Surprisingly, *Danio rerio* GEMIN3 did not form a ternary complex with VHL despite 80% conservation of the zebrafish sequence. Chimeric human-zebrafish constructs were designed to comprehensively cover all sequence differences between the orthologs and were tested by NanoBiT for their effect on ternary complex formation. Four constructs representing two regions, GEMIN3_aa197-220_ and GEMIN3_aa236-260_, imbued the chimeras with partial loss-of-function when inserting zebrafish sequences into human GEMIN3, and partial gain-of-function when inserting human sequences into zebrafish GEMIN3 (Fig. 5b). We rationalized that any degron recognized by VHL would be on the surface of GEMIN3 and designed additional chimeras to test solvent-exposed sequences in the two regions of interest. Two shorter sequences, GEMIN3_aa219-220_ and GEMIN3_aa249-256_, reproduced the partial loss- or gain-of-function seen with the larger constructs (Fig 5c).

**Fig. 5.**
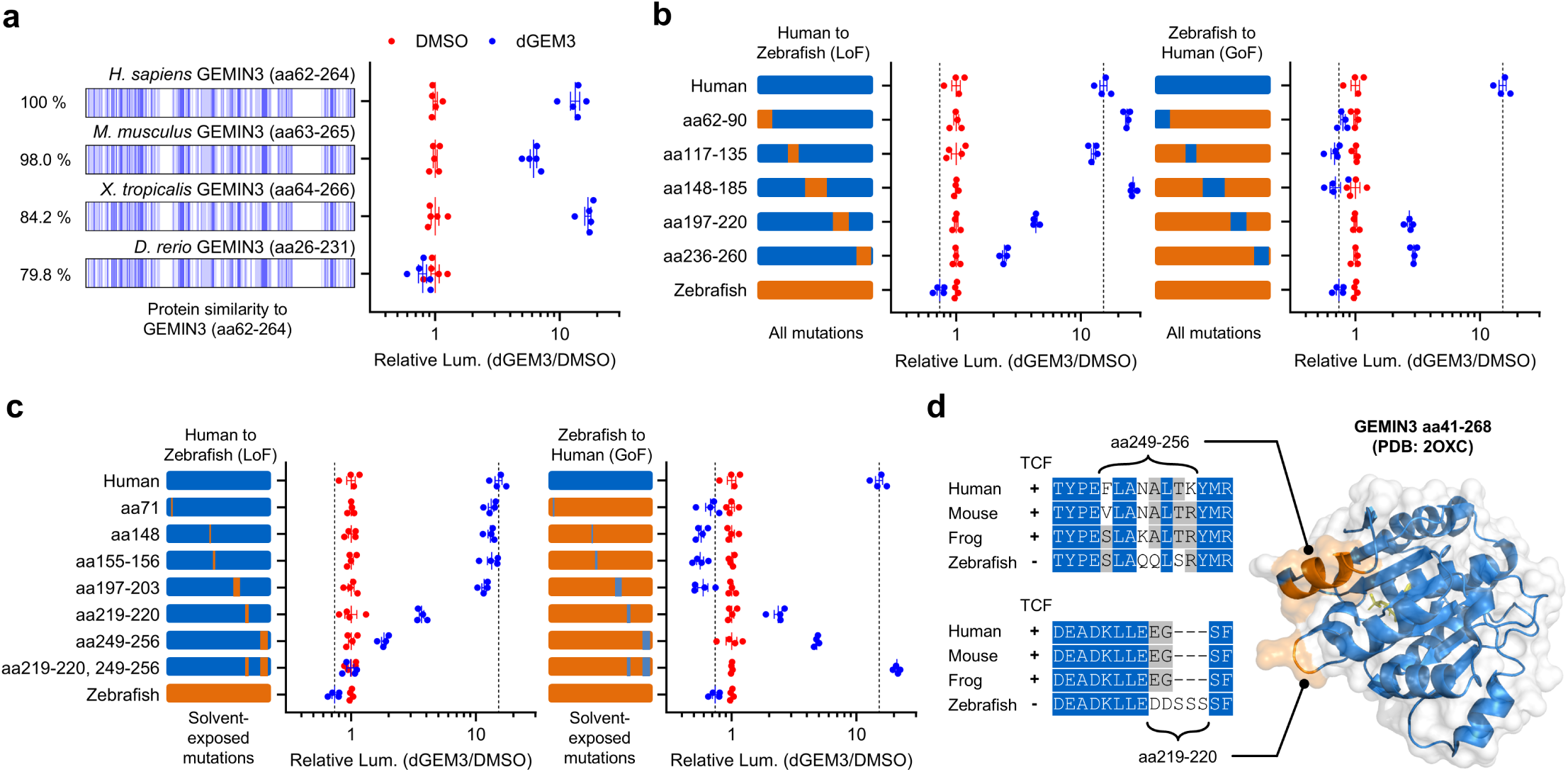
Mapping the GEMIN3 degron based on ortholog-selectivity of dGEM3. **a**, Selectivity for GEMIN3 homologs by VHL ternary complex assay. Helicase ATP-binding domains alignment shown as in Fig. 1f. Cells pre-treated with BTZ were subjected to 10 μM dGEM3 or DMSO treatment for 2 h and luminescence was calculated per-construct relative to DMSO controls. Mean ± s.e.m.; n = 5. **b**, TCF of Zebrafish-Human GEMIN3 chimera constructs. Constructs were designed to account for all mutations between human (blue) and zebrafish (orange) sequences in the GEMIN3 helicase ATP-binding domain. Cells expressing VHL and the chimeric GEMIN3 constructs were treated with 10 μM dGEM3 for 2 h to monitor TCF. Dashed lines correspond to WT human GEMIN3 positive control and WT zebrafish GEMIN3 negative control. Mean ± s.e.m.; n = 4. **c**, TCF of solvent-exposed Zebrafish-Human GEMIN3 mutations. Same as Fig. 5b but only solvent-exposed residues are mutated. Cells expressing VHL and the GEMIN3 mutant constructs were treated with 10 μM dGEM3 for 2 h to monitor TCF. **d**, Human GEMIN3 degron mapped to corresponding crystal structure (PDB: 2OXC) and alignment of homolog sequences necessary for TCF. The GEMIN3_aa219-220_ loop and GEMIN3_aa249-256_ helix are shaded in orange. See Fig. S3 for full alignment.

Gratifyingly, mutating these two solvent-exposed sequences together fully abolished or reinstated TCF in their respective chimeras, proving that GEMIN3_aa219-220_ and GEMIN3_aa249-256_ are both necessary and sufficient for dGEM3-induced TCF between VHL and GEMIN3. Alignment of the ortholog sequences for these regions reveals bulky substitutions at A253Q and G220D, as well as the insertion 220-221insSSS in zebrafish GEMIN3 that could produce a steric clash during TCF (Fig. 5d). When mapped to the GEMIN3 crystal structure, these two sequences identify a loop and helix that are directly adjacent and likely make up a composite structural degron recognized by VHL.

### Impact of dGEM3 SAR on ternary complex kinetics and degradation

To gain insight into the structural basis of the molecular glue interaction, we modeled the VHL:dGEM3:GEMIN3_aa41-268_ ternary complex using *in silico* protein-protein docking. By constraining the model based on the predicted GEMIN3 degron and calculating scores for the resulting interfaces, we consistently observed that high-scoring models made close contact between GEMIN3 and the dichlorobenzene of dGEM3. This observation inspired the rational design of several dGEM3 analogs, compounds **3** through **6** (Fig. 6a). We systematically excluded the RHS dichlorobenzene ortho and para chlorine atoms to assess their contribution to VHL binding, ternary complex formation, and GEMIN3 degradation. Additionally, the oxygen atom in the left-hand side oxazinane ring was replaced with a carbon atom, forming a fused pyrrazole, to evaluate the contribution from the LHS of the molecular glue.

**Fig. 6.**
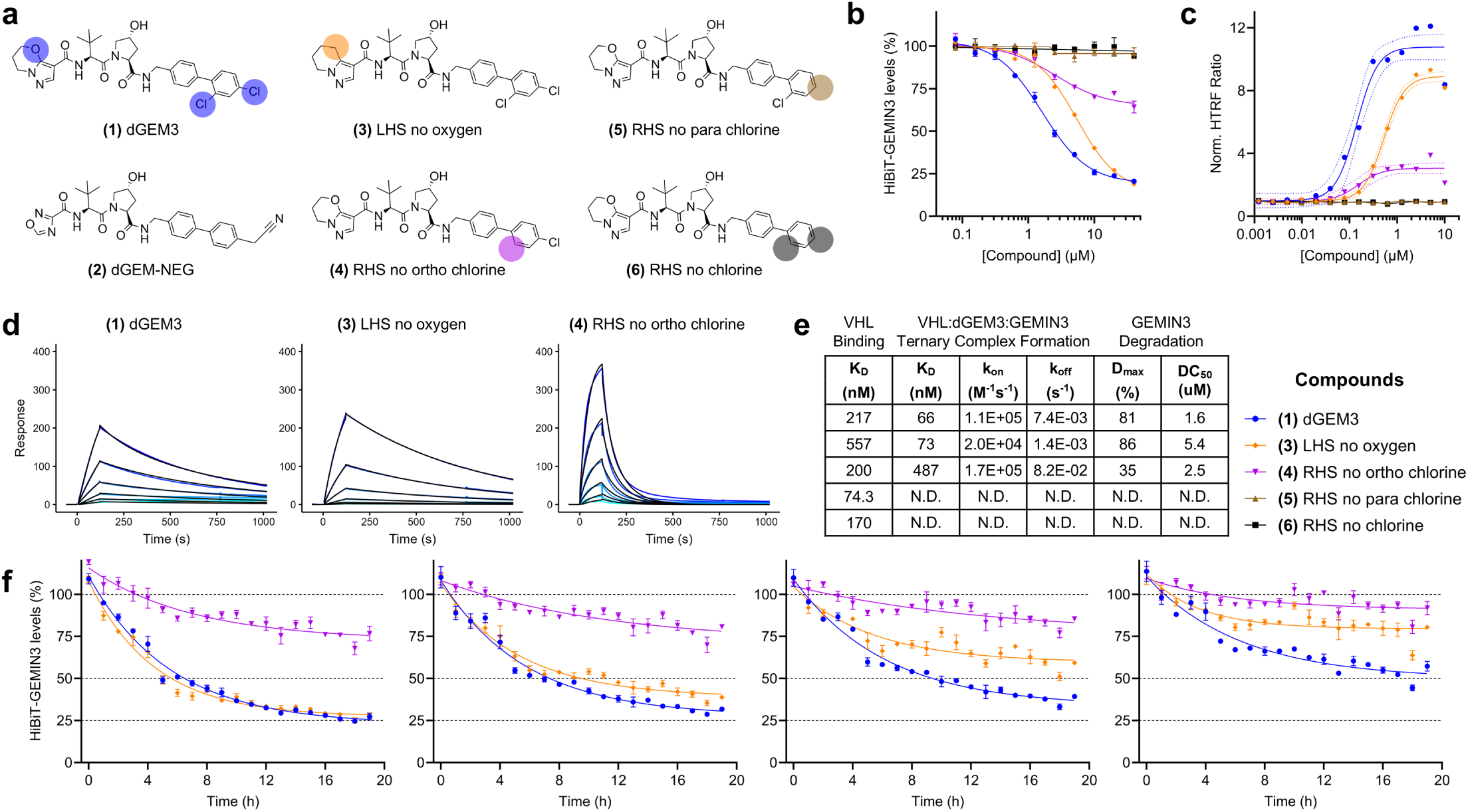
Impact of dGEM3 SAR on ternary complex kinetics and degradation. **a**, Chemical structures of dGEM3 analogs. Differences among the analogs are shaded by color and match subsequent panels. **b**, Cellular GEMIN3 degradation by dGEM3 analogs. HiBiT-GEMIN3 HEK293T cells were treated with compounds in dose-response up to 40 μM for 15 h. Mean ± s.e.m.; n = 3. **c**, *In vitro* VHL TCF with dGEM3 analogs. VHL was incubated with GEMIN3_aa41-268_ and compound for 1 h as in Fig. 4a. Solid lines indicate regression curve fit to 14-point 2-fold dose-response, and dashed lines indicate 95% confidence bands for nonlinear fit. **d**, VHL:dGEM3:GEMIN3_aa41-268_ ternary complex formation by SPR. The VBC:compound complex was pre-formed and analyzed for TCF by multi-cycle kinetics on a GEMIN3-functionalized surface with drug concentrations of 25, 50, 100, 200, and 400 μM. Kinetic parameters listed in Fig. 6e. **e**, Summary of parameters for VHL binding, ternary complex formation, and GEMIN3 degradation by dGEM3 analogs. K_D_, equilibrium dissociation constant, k_on_, association rate constant; k_off_, dissociation rate constant; D_max_, maximal degradation; DC_50_ half-maximal degradation concentration. **f**, Time-course of cellular GEMIN3 degradation by dGEM3 analogs. HiBiT-GEMIN3 cells were treated with varying doses of active dGEM3 analogs for 19 h. Solid lines are best fit curves (one-phase decay) of means from n=3 independent treatments with s.e.m. represented by error bars.

Measuring GEMIN3 degradation by the dGEM3 analogs, the narrow structure activity relationship (SAR) for the molecular glue became immediately apparent. While removal of the RHS ortho chlorine in compound **4** profoundly decreased the D_max_ of the compound, removing the para chlorine alone in compound **5** completely abolished degradation of GEMIN3 (Fig. 6b). Modification to the LHS of dGEM3 by removing the oxygen in compound **3** decreased the potency but preserved the same maximal degradation at higher doses. To investigate whether changes in ternary complex formation were responsible for the impaired degradation activity of the analogs, dGEM3 and compounds **3** through **6** were tested *via* the established *in vitro* TCF HTRF assay. Consistent with the degradation assay, removing the para chlorine of the dGEM3 RHS dichlorobenzene alone (compound **5**) or in combination with the ortho chlorine (compound **6**) completely abolished TCF (Fig. 6c). Removal of the LHS oxygen or RHS ortho chlorine from dGEM3 preserved some level of TCF activity in compounds **3** and **4** respectively, but increased the K_D_ for the ternary complex. As compounds **5** and **6** demonstrated no activity in the degradation or TCF assays, we used SPR to confirm that they were still binding to VHL (Fig. 6e, S4a). Indeed, all the analogs bound VHL with binary K_D_ values similar to VH298, confirming that the loss of GEMIN3 degradation was caused by loss of the ternary complex.

Having established absolute dependence of dGEM3 on the RHS para chlorine for molecular glue activity, we sought to fully characterize the remaining analogs with impaired potency, compounds **3** and **4**. We used SPR to examine the ternary complex kinetics for dGEM3 analogs modified on the LHS (no oxygen) or RHS (no ortho chlorine). To our surprise, modifying the LHS and RHS of dGEM3 separately impacted the association rate (k_on_) and dissociation rate (k_off_), with the LHS no oxygen analog (compound **3**) impaired by slower k_on_ and the RHS no ortho chlorine analog (compound **4**) impaired by faster k_off_ (Fig. 6d, 6e, S4b). Of the two, k_off_ appears to have a far larger impact on GEMIN3 degradation, with the fast k_off_ analog demonstrating a dramatically impaired D_max_ relative to dGEM3 (35% versus 81%) (Fig. 6e, 6f). This effect on D_max_ was corroborated by BioE3 proteomics, which showed reduced GEMIN3 ubiquitination in line with the shorter-lived ternary complex formed by compound **4** (Fig. S4c). To test whether the relative impairments to ternary complex k_on_ and k_off_ could be overcome for effective targeted protein degradation, compounds **3** and **4** were tested at increasing time points and doses. While slow ternary complex association could be overcome by increasing time and dose, neither could compensate for rapid ternary complex dissociation (Fig. 6f). This set of analogs confirm the importance of optimizing kinetics for long-lived ternary complexes not only for PROTAC development, but also in the development of molecular glues for the VHL E3 ligase.

## Discussion

VHL has been the subject of research for more than 30 years—first as a tumor suppressor associated with heritable cancers, second as a key E3 ligase endogenously regulating hypoxia, and most recently as a reliable E3 ligase effector for PROTAC development. This study highlights and corroborates an emerging role for VHL as a “glue-able” E3 ligase amenable to redirecting by monovalent molecular glues.

VHL molecular glues have evaded detection for several reasons, among them a dearth of screening methodologies specifically tailored to molecular glues. While this has recently begun to be addressed by various target-agnostic approaches^11^, it was anticipated that the Picowell RNA-seq approach combining a similar unbiased readout with a biased small molecule library design would be key to the discovery of novel glues. In the design of a VHL molecular glue library with tunable specific interactions, a diverse set of exit vectors that included various structural modifications at multiple positions on the core binding scaffold were incorporated. This strategy aimed to include diverse functional groups to facilitate the effective engagement of the surface of VHL with potential neosubstrates. While DELs have previously been used to explore VHL PROTACs^67^, we believe this is the first application of a VHL-focused DEL in a cell-based phenotypic screen.

dGEM3 demonstrated a remarkable degree of selectivity for GEMIN3 recruitment directly out of the DEL screen with no other DEAD-box helicases showing any indication of recruitment or degradation. The SAR around dGEM3 was similarly narrow, with GEMIN3 binding to VHL dependent on a single chlorine atom in dGEM3. These observations suggest VHL is capable of highly selective TPD but may also be subject to inflexibility in the protein-protein interface between VHL and GEMIN3. Adding intrigue, the VHL inhibitor VH032 has been reported to possess molecular glue activity against cysteine dioxygenase 1 (CDO1), which has an entirely unrelated protein fold to GEMIN3. Combined with the knowledge that VHL is a versatile degrader of diverse targets using PROTACs, the ability of VHL to target additional proteins using molecular glues remains an open question. We anticipate that elucidation of the VHL:dGEM3:GEMIN3 structure should provide significant insight into any potential shared degrons among VHL neosubstrates not readily apparent by sequence or conserved fold.

The recruitment of GEMIN3 to VHL by dGEM3 exemplifies TCF, ubiquitination, and degradation as key principles of TPD shared by molecular glues and PROTACs alike. Our multi-omics profiling of dGEM3 by TurboID, BioE3, and global proteomics (Fig. 2b-d) provides a comprehensive characterization of the MGD’s activity, both rationalizing the exquisite selectivity for GEMIN3, and highlighting the necessity of each step in the process of protein degradation. Previous PROTACs have demonstrated collateral degradation of indirectly recruited members of HDAC complexes^68^, and DDB1 MGDs instead degrade only the indirect cyclin K interaction partner of their CDK12 target^26,29^. Our data confirm the possibility of observing glue-induced proximity and even ubiquitination of proteins, such as SMN2, that does not lead to their effective degradation. Considering additional recent insights from proteomics methods demonstrating complex molecular glue and PROTAC polypharmacology against targets that are spared from degradation^69,70^, there is a strong rationale for considering ternary complex formation, ubiquitination, and degradation separately in the context of degrader discovery and optimization.

The requirement for ARIH1 and UBE2L3 as “priming” ligases for *in vitro* ubiquitination of GEMIN3 by VHL also serves as a reminder that factors aside from E3 ligase and target play key roles in the efficiency of TPD. Functional screens for genetic dependencies of degraders^71^ and UPS components^72^, as well as proteomic profiling of dynamic E3 ligase assembly^73,74^, each highlight the heterogeneity of E3 ligase activity and regulation in the UPS. While this study benefited from extensive work by other researchers establishing regulatory mechanisms for CUL2 ligases like VHL, we anticipate that future molecular glue discovery efforts will require functional genomics support as the E3 ligase toolbox expands.

Lastly, SAR explored in the study provides reassurance that optimizing stable ternary complex formation is a worthwhile path for degrader optimization. Although modulating ternary complex kinetics has been demonstrated using PROTACs^66,70,75,76^, we are not aware of a molecular glue example demonstrating the influence of ternary complex dissociation kinetics on degradation. Though unanticipated, the residual activity of the para chlorine-containing dGEM3 analogs provided a convenient way to separately test these kinetic parameters. The molecular glues profiled here concretely show that prolonging sufficient ternary complex half-life through slow dissociation is necessary to achieve maximal protein degradation. Although this conclusion is consistent with expectations of E3 ligase function, the limited number of MGD examples has precluded confirmation of this hypothesis until now.

In summary, our research provides an example of rational molecular glue discovery for VHL, identifying dGEM3 as a molecular glue degrader targeting the GEMIN3 helicase ATP-binding domain. We anticipate that unbiased phenotypic screening in combination with biased E3-focused small molecule libraries will continue to be a fruitful approach to molecular glue discovery. Finally, our discovery of GEMIN3 as a VHL neosubstrate expands the known molecular glue target space of VHL, furthering the possibility of redirecting VHL to degrade new and more therapeutically relevant targets.

## Supplemental Figures

**Fig. S1.**
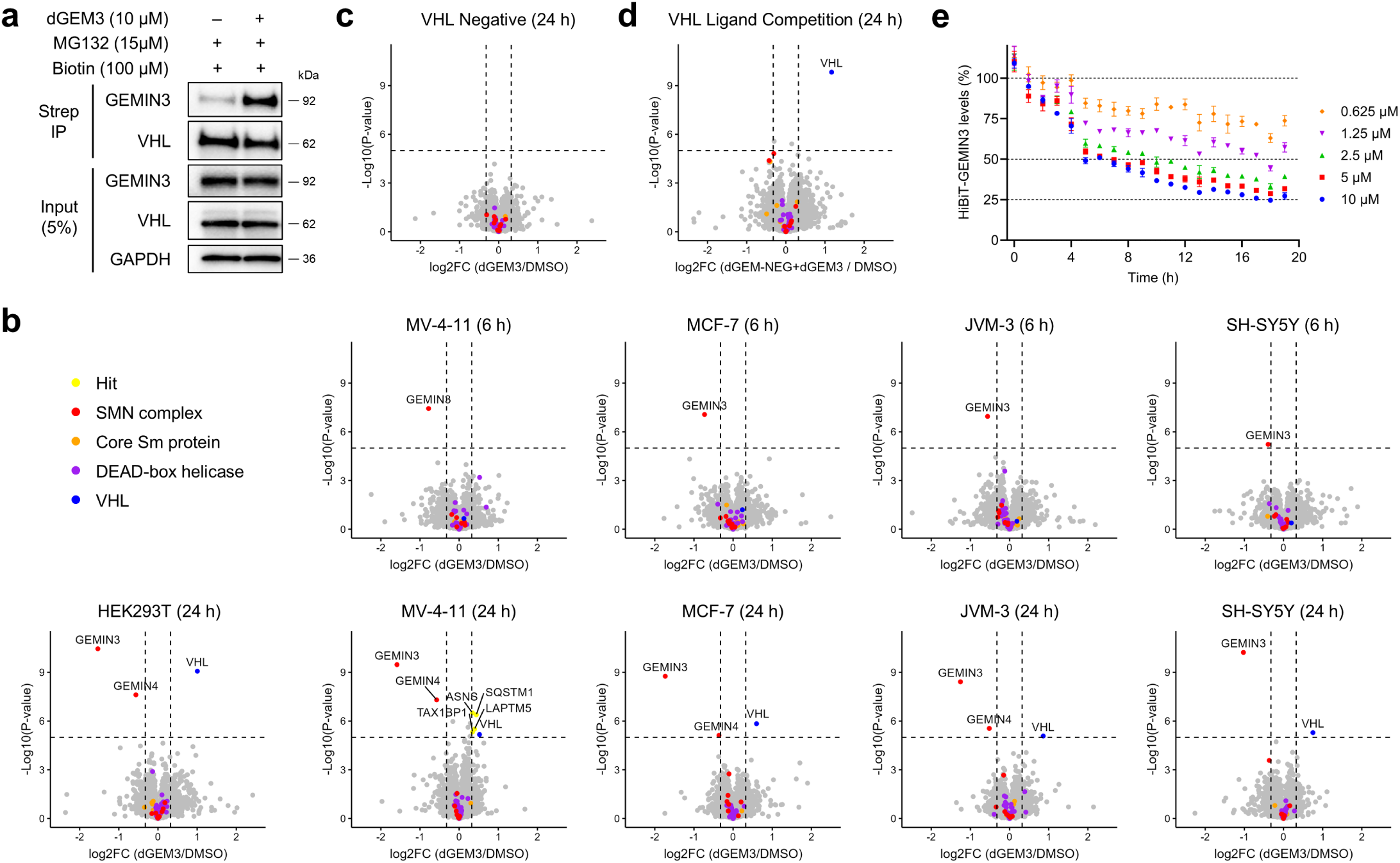
Extended characterization of GEMIN3 degradation by dGEM3. **a**, Immunoblot of TurboID-VHL proximity labeling assay from Fig. 2b for GEMIN3 and VHL enrichment. Representative of three biological replicates. **b**, Global proteomics following treatment with 10 μM dGEM3 (n=4) or 0.1% DMSO (n=4) in HEK293T, MV-4-11, MCF-7, JVM-3, and SH-SY5Y cancer cell lines cell lines at 6 and 24 h time points. Dashed lines indicate significance and enrichment cutoffs of p < 0.00001 and FC > 1.25. Legend same as Fig. 2d. **c**, Global proteomics of 24 h 10 μM dGEM3 treatment in VHL-deficient 786-O cancer cell line. Replicates, cutoffs, and colors are identical to Fig. 2d. **d**, Global proteomics of 24 h 10 μM dGEM3 treatment following pre-treatment with competitor dGEM-NEG. Replicates, cutoffs, and colors are identical to Fig. 2d. **e**, GEMIN3 cellular degradation assay in endogenously tagged HiBiT-GEMIN3 HEK293T cell line. Cells were treated in 19 h time-course with dGEM3 concentrations up to 10 μM. Dashed lines indicate 0%, 50%, and 75% degradation. Mean ± s.e.m.; n = 3 independent treatments per dose and time point.

**Fig. S2.**
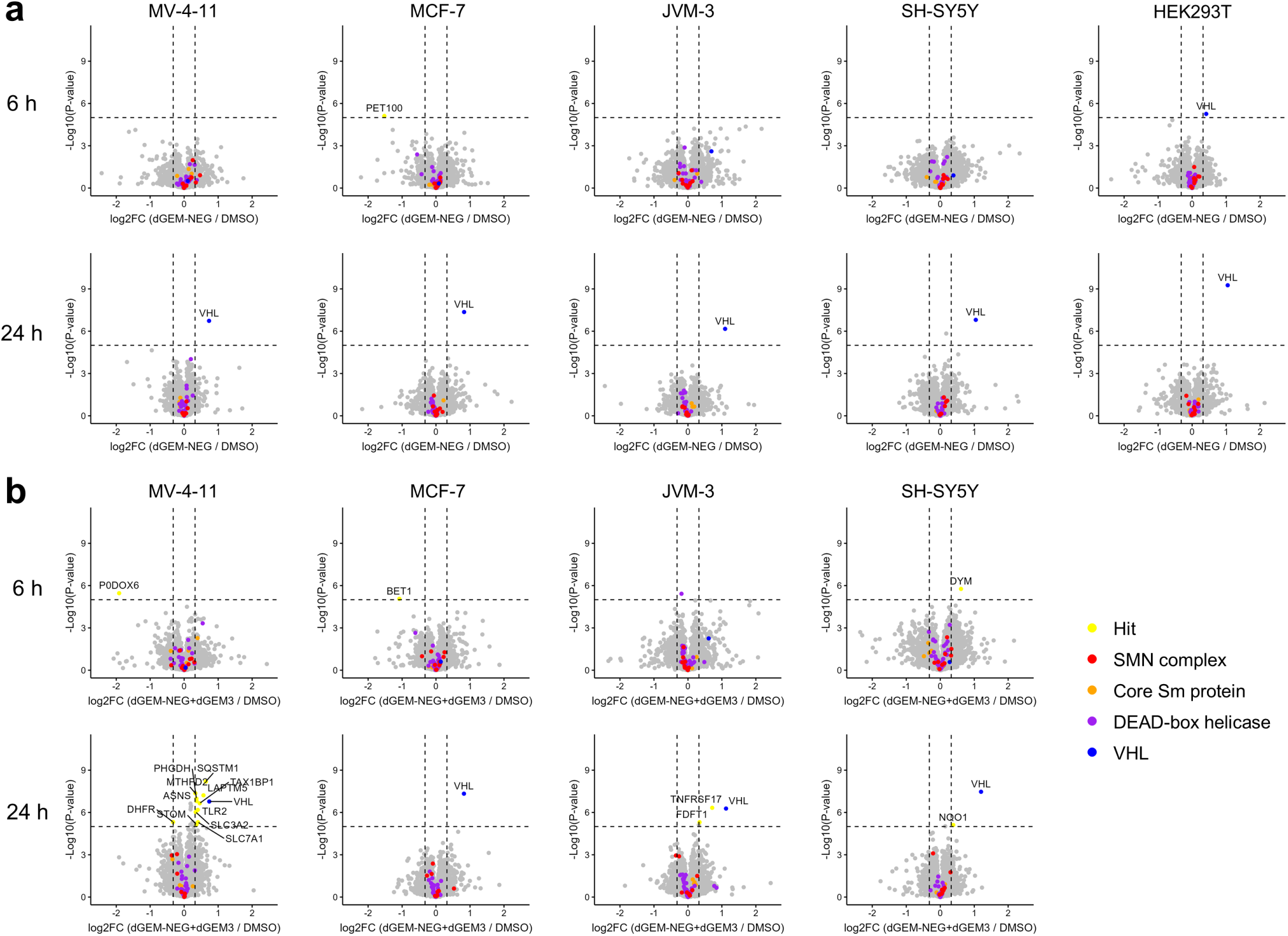
Extended characterization of dGEM-NEG activity. **a**, Global proteomics of negative control compound dGEM-NEG. MV-4-11, MCF-7, JVM-3, SH-SY5Y, and HEK293T cancer cell lines were treated with 10 μM dGEM-NEG (n=4) or 0.1% DMSO (n=4) for 6 or 24 h. Replicates, cutoffs, and colors are identical to Fig. 2d. **b**, Global proteomics of dGEM3 treatment with dGEM-NEG VHL ligand competition. Cells were pre-treated with 10 μM dGEM-NEG for 0.5 h before addition of 10 μM dGEM3. Replicates, cutoffs, and colors are identical to Fig. 2d.

**Fig. S3.**
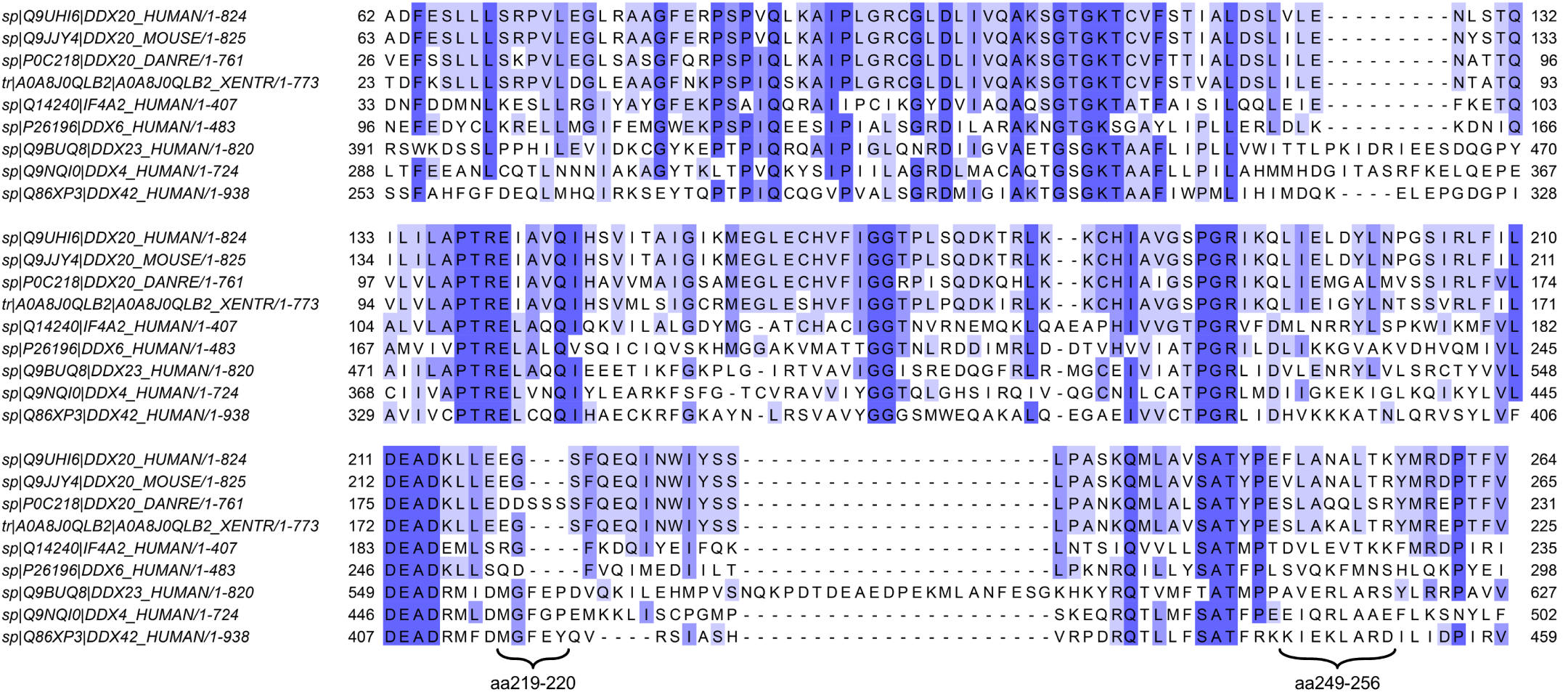
Sequence alignment of tested constructs. **Clustal Omega multiple** sequence alignment (MSA) of GEMIN3, GEMIN3 homologs, and members of the DEAD-box helicase family. Sequences correspond to helicase ATP-binding domain constructs used for cellular NanoBiT assays. Mutated candidate degron from Fig. 5d indicated by brackets.

**Fig. S4.**
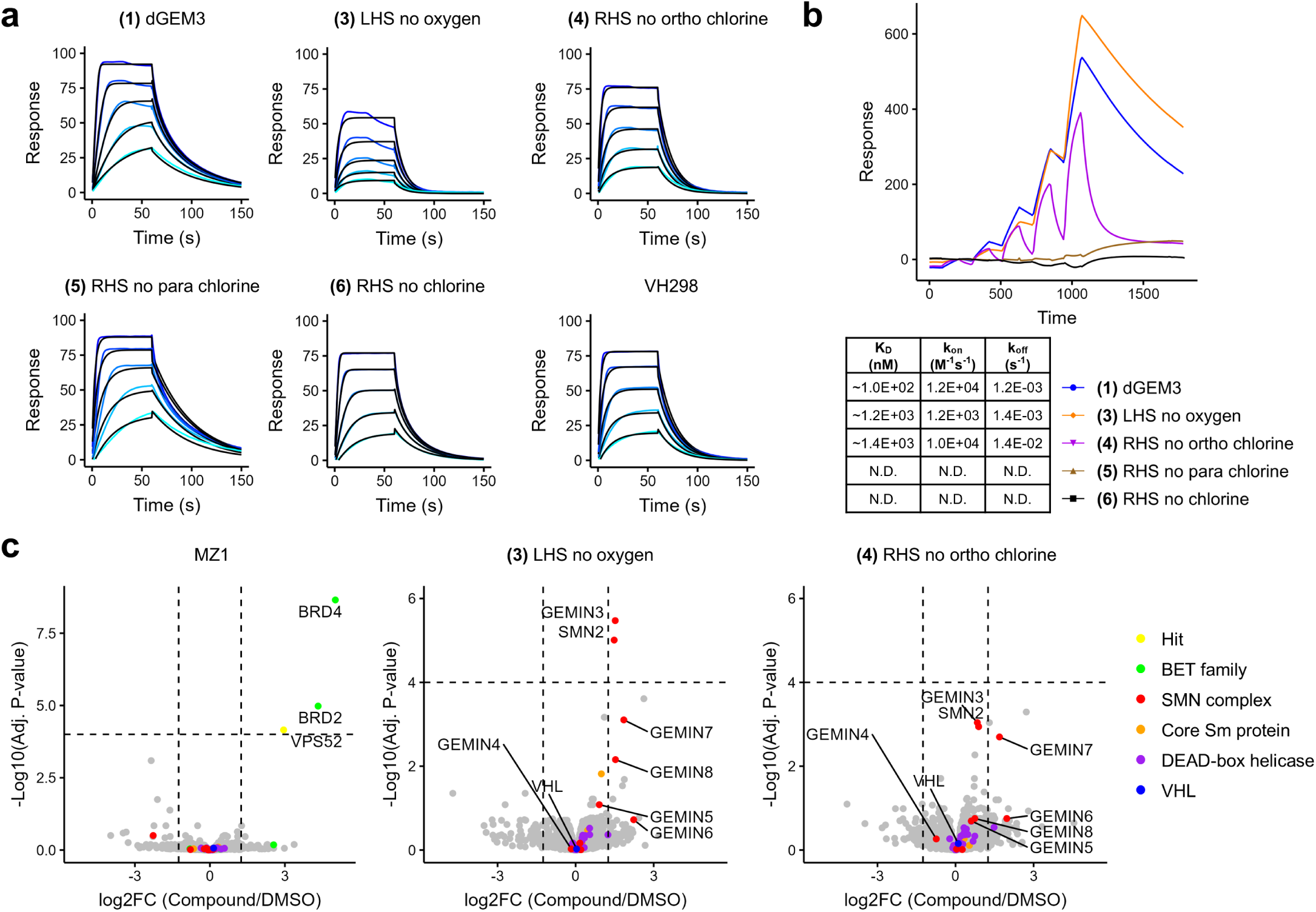
Impact of dGEM3 SAR on VHL binding and ternary complex formation. **a**, Binary VHL-binding of dGEM3 analogs by SPR. Compounds were analyzed in kinetic multi-cycle format on a VBC-functionalized chip. K_D_ values are listed in Fig. 6e. **b**, VHL:dGEM3:GEMIN3_aa41-268_ ternary complex formation by SPR. The VBC:compound complex was pre-formed and analyzed for TCF by single-cycle kinetics on a GEMIN3-functionalized surface with drug concentrations of 25, 50, 100, 200, and 400 μM. **c**, BioE3 VHL proximity ubiquitin labeling assay following 2 h co-treatment with 2 μM BTZ and 10 μM MZ1, compound 3, or compound 4 (n=3) compared against 0.1% DMSO (n=3). Significance, enrichment cutoffs, and legend the same as Fig. 2c. BET family proteins indicated in green for positive control MZ1. BTZ, bortezomib.

## Materials and Methods

### Cell Culture

HEK293 (CRL-1573), HEK293T (CRL-3216), 786-O (CRL-1932), K-562 (CCL-243), MV-4-11 (CRL-9591), MCF-7 (HTB-22), and SH-SY5Y (CRL-2266) cells were obtained from ATCC, and JVM-3 (ACC 18) was obtained from DSMZ. HEK293, HEK293T and MCF7 cells were cultured in DMEM, JVM-3 and 786-O were cultured in RPMI, K-562 and MV-4-11 were cultured in IMDM, and SH-SY5Y was cultured in advanced DMEM/F12 medium. All media was supplemented with 10% FBS, 50 U/mL penicillin and 50 U/mL streptomycin. Cells were grown at 37°C in a 5% CO_2_ incubator and underwent regular testing for mycoplasma contamination.

### Picowell RNA-seq Screen

Solid phase synthesis was used to produce beads containing compounds and DNA oligos which encode the compound identity. Oligos also contain a poly-T stretch to capture poly-adenylated RNAs from lysed cells. Bead-based libraries were loaded on custom microfluidic devices (containing ∼56,000 picowells) with one bead per picowell. Cells were loaded achieving an average density of 40 cells per picowell, then covered with a proprietary isolation solution to prevent compound leakage. Cells were incubated for 2 hours at 37°C before compounds were photocleaved from beads using UV light to achieve an effective concentration of ∼10 μM in each well. Microfluidic devices containing compound-treated cells were incubated for 24 hours at 37°C. Without breaking isolation, cells were lysed, and poly-adenylated RNA was captured on the beads by the poly-T containing oligos. Bead-RNA complexes were removed from the microfluidic devices and washed using high-salt buffers with RNase inhibitors to remove cell debris but retain hybridized RNA. On-bead RNA was subjected to reverse transcription and cDNA followed by RNA-seq library construction. Picowell RNA-seq libraries were then sequenced on an Illumina Novaseq system. Analyses was performed utilizing custom scripts in R Statistical Software^77^. Significant gene signatures and compounds were identified, and compounds were subsequently resynthesized off-bead for hit validation.

### VHL NanoBRET Target Engagement Assay

VHL binding affinity was assessed using the VHL NanoBRET Target Engagement Intracellular E3 Ligase Assay (Promega #N2930) in live-cell assay mode following the manufacturer protocol. In brief, HEK293T cells were transfected in bulk with NanoLuc-VHL expression plasmid using Lipofectamine 3000 (Invitrogen #L3000015). The next day, cells were replated in Opti-MEM in 384-well format (2 × 10^5^ cells/mL, 38 μL/well) and VHL Tracer Reagent was added (2 μL, 1 μM final concentration). Compounds were added directly to wells using a Tecan D300e digital dispenser and normalized to 0.5% DMSO. Compound-treated cells were incubated for 2 hours then equilibrated to room temperature. NanoBRET Nano-Glo substrate and extracellular NanoLuc Inhibitor was freshly prepared in Opti-MEM at 3X concentration and dispensed 20 μL/well. Following quick spin and mixing, plate was incubated on shaker for 3 minutes. The assay was read on a BMG CLARIOstar multimode plate reader using 450 nm donor emission and 610 nm acceptor emission. Data were analyzed using GraphPad Prism software, normalizing to the max and min across the plate and fitting to a four-parameter logistic curve.

### Bulk RNA-seq

HEK293T cells were treated with 10 μM compound or 0.1% DMSO control for 6 hours. RNA was isolated using the miRNeasy Mini kit (Qiagen #217004) with on-column DNAse digestion using the RNase-Free DNase set (Qiagen #79254). RNA integrity number (RIN) was confirmed to be >8 using an Agilent 2100 Bioanalyzer. RNA-seq libraries were prepared using 500 ng of total RNA and the QuantSeq 3’ mRNA-Seq V2 Library Prep Kit (Lexogen) in 96-well plates. All 96 libraries were pooled and sequenced in an Illumina Nextseq550 to obtain approximately 5 million single-end 75 bp reads per sample. Adapters, poly-A tails, and low-quality bases were removed using Cutadapt^78^, reads mapped to the human genome (hg38) using STAR^79^, raw counts obtained using featureCounts^80^, and differential gene expression analysis carried out with DESeq2^81^.

### TurboID Cell Treatments

K-562 cells stably overexpressing TurboID-VHL were plated in 10 cm dishes at 0.5 × 10^6^ cells/mL and cultured overnight. The next day, cells were pre-treated with 15 µM MG-132 (Selleck Chemicals #S2619) for 1 hour. Next, 10 µM dGEM3 and 100 µM D-Biotin (Invitrogen #B1595) were added and cells were incubated for 6 hours. To harvest, cells were gently washed three times with 10 mL of ice-cold PBS, then resuspended in 300 µL RIPA buffer (Thermo Scientific #89900) containing cOmplete Protease Inhibitor Cocktail (Roche #4693116001). After 20 minutes, lysate was clarified by high-speed centrifugation (14,000 RPM, 10 minutes) and 250 µl was subjected to streptavidin pull down using 40 µL of pre-washed Pierce Streptavidin Magnetic Beads (Thermo Scientific #88817) brought up to 500 µL final volume in RIPA buffer. Sample tubes were rotated overnight at 4°C before washing by rotating at room temperature, twice in 1 mL RIPA buffer for 10 minutes, once in 1 mL Pierce IP Lysis buffer (Thermo Scientific #87787) for 10 minutes, and once in 1 mL PBS for 1 hour.

For western blot analysis, half of the beads were mixed with 50 µL 2X SDS sample loading buffer with 5 mM D-Biotin and 50 mM DTT, then incubated with shaking at 95°C for 10 minutes. Eluted samples were subject to western blot and probed using primary antibodies for GEMIN3 (Proteintech #11324-1-AP), VHL (Cell Signaling #68547), and GAPDH (Cell Signaling #2118).

### TurboID Sample Preparation for LC-MS Analysis

For LC-MS analysis, each streptavidin pull-down bead sample generated from the TurboID assay was reconstituted in 40 µL of 50 mM Tetraethylammonium Bromide (TEAB) buffer. Reduction and alkylation were performed by adding 50 mM chloroacetamide (CAA) and 10 mM Tris(2-carboxyethyl)phosphine (TCEP) to each sample, followed by incubation at 56°C for 45 minutes to denature and stabilize proteins. For on-bead digestion, 1.5 µg of a Trypsin/Lys-C mixture was added to each sample, and the reaction was incubated at 37°C overnight. Following digestion, the samples were transferred to new tubes and subjected to a single-pot, solid-phase-enhanced sample preparation (SP3) peptide cleanup protocol. Briefly, 10 µL of SP3 beads were added to each sample, and peptides were bound to the beads by adjusting the acetonitrile concentration to 95% (v/v) and incubating at room temperature for 10 minutes. Beads were then washed three times with 500 µL of 100% acetonitrile to remove contaminants. Finally, peptides were eluted from the beads in 50 µL of MS-grade water and dried using a SpeedVac concentrator.

### TurboID LC-MS Data Acquisition

Dried samples were reconstituted in 0.1% FA prior to injection. Samples were then analyzed on a Vanquish-Neo (Thermo Fisher Scientific) Orbitrap Eclipse, LC-MS system. Detailed acquisition methods are as follows: Samples were separated using a 15-cm PepSep C18 column at a flow rate of 1.2 µL/min. A multi-step gradient was applied using mobile phase B (80% acetonitrile with 0.1% formic acid). Initially, the B-phase was increased from 1% to 2% over the first 0.1 minutes. This was followed by a linear ramp from 2% to 24% between 0.1 and 45.1 minutes, then a further increase from 24% to 40% from 45.1 to 60.1 minutes. Finally, the gradient was held at 95% from 60.1 to 70.6 minutes to ensure effective elution of strongly retained components and proper column re-equilibration. For mass spectrometry acquisition, the ion transfer tube was maintained at 300 °C and a voltage of 2200 V was applied to the emitter to ensure efficient peptide ionization. Data-dependent acquisition (DDA) mode was used. Full MS1 scans were recorded in the Orbitrap at a resolution of 60,000, with the RF lens set to 40% and an AGC target of 8×10^5 ions. Peptide precursor ions bearing charge states from 2+ to 7+ were selected and isolated in the quadrupole with a 0.7 Th isolation window. These precursors were then subjected to higher-energy collisional dissociation (HCD) at a normalized collision energy of 33%, and the resulting MS2 spectra were acquired in the Orbitrap at a resolution of 15,000 with an AGC target of 5×10^4 ions.

### TurboID LC-MS Data Processing and Analysis

Raw data were processed using FragPipe (v18) with the DDA-LFQ workflow. MS/MS spectra were searched against the SwissProt Human canonical reference proteome database using the following parameters: peptides were restricted to a length of 7–50 amino acids, and enzyme specificity was set to restricted trypsin digestion (cleaving after lysine and arginine). Carbamidomethylation of cysteine was defined as a fixed modification, while N-terminal acetylation and methionine oxidation were allowed as variable modifications. FDR was controlled at 1% at both the peptide and protein levels using Peptide Prophet and Protein Prophet^82^. Protein quantification was performed using IonQuant^83^. Subsequent statistical analysis was carried out utilizing custom scripts in R Statistical Software^77^ that normalized protein quantification results to VHL and applied statistical modeling and hypothesis testing using a two-sided moderated t-test as implemented in the limma package^84^.

### BioE3 Cell Treatments

HEK293 cells were engineered by knockout of endogenous VHL using CRISPR, overexpression of VHL-BirA using the PiggyBac transposon system, and introduction of a Tet-On bioGEF-Ub_L73P_ expression cassette at the AAVS1 safe harbor locus^53,85^. For BioE3 cell treatments, cells were plated in DMEM + 10% dialyzed FBS, 1% Pen-Strep, and 1mM sodium pyruvate in 96-well format with 2.5 × 10^4^ cells per well. Following 24-hour treatment with 0.5 µg/mL doxycycline, cells were co-treated with 10 µM dGEM3 and 2 µM bortezomib for 2 hours. Next, biotin was added to the wells at a final concentration of 50 µM for 30 minutes. Cells were then washed with PBS four times and stored at -80°C until further processing for mass spectrometry.

### BioE3 Sample Streptavidin Enrichment and LC-MS Sample Preparation

Cells cultured and treated in a 96-well plate were lysed in 150 µL of RIPA buffer supplemented with 25 units of BenzoNuclease and a 1x Protease Inhibitor Cocktail. The cell lysates were subjected to sonication at 50 Hz for 20 minutes using a PIXUL sonicator (Active Motif) to ensure efficient cell disruption. For enrichment, 15 µL of high-capacity streptavidin magnetic beads were added to the lysates, and the mixture was incubated to allow binding. Following incubation, the beads were sequentially washed six times with various buffers to remove non-specifically bound proteins: twice with RIPA buffer, once with 8 M Urea, once with 5 M NaCl, and twice with 50 mM EPPS buffer. On-bead digestion was carried out by adding 0.5 µg of a Trypsin/Lys-C mixture in 50 µL of 50 mM EPPS buffer per sample and incubating overnight at 37°C. Subsequently, 40% of each digested sample was loaded onto an Evotip for downstream analysis.

### BioE3 LC-MS Data Processing and Analysis

Samples were analyzed on an EvoSep TIMSTOF-HT LC–MS system. Peptide separations were performed using the Evosep One 100SPD method (i.e. throughput of 100 samples per day) on an 8-cm PepSep C18 analytical column. The TIMSTOF-HT operated in diaPASEF mode, with ion mobility separation spanning an inverse reduced mobility (1/K₀) range of 0.7–1.3 V·s/cm², mass range from 100-1700 m/z. Both the ramp time and accumulation time were set to 65 ms, and 36 scan windows were acquired per cycle, yielding a total cycle time of 0.85 s. diaPASEF data were searched against an in-house built human spectral library using DIA-NN (v1.8.1), protein quant results were analyzed through a custom R script that normalized protein quantification results to VHL and applied statistical modeling and hypothesis testing using the limma package^77,84,86,87^.

### Global Proteomics Cell Treatments

For global proteomic experiments, HEK293T, MV-4-11, MCF-7, JVM-3, SH-SY5Y, and 786-O cell lines were plated in 96-well format and grown overnight. The next day, compounds were added to each well at 10 μM concentration and incubated for 6 or 24 hours. In competition experiments, cells were pre-treated with 10 μM dGEM-NEG before addition of dGEM3. Following compound treatments, cells were washed in the 96-well plate three times with PBS and centrifuged before lysis.

### Global Proteomics Sample Preparation

Sample preparation for global proteomics was performed in 96-well plates. First, cells were lysed by adding lysis buffer (8M urea, 50 mM ammonium bicarbonate, and Benzonase 24U/100mL); and after sealing, the plate was subjected to vigorous shaking in a Retsch MM301 instrument (20 Hz for 10 min at room temperature). Protein concentration of the supernatant was determined using bicinchoninic acid (BCA) protein assay (Thermo Scientific #23227). Proteins were reduced with 5 mM tris(2-carboxyethyl)phosphine (TCEP) at 30°C for 60 min, and subsequently alkylated with 15 mM iodoacetamide (IAA) in the dark at room temperature for 30 min. Urea was then diluted to 1 M urea using 50 mM ammonium bicarbonate, and proteins were subjected to overnight digestion with mass spec grade Trypsin/Lys-C mix (1:25 enzyme/substrate ratio). Following digestion, samples were acidified with formic acid (FA) and subsequently desalted using AssayMap C18 cartridges mounted on an Agilent AssayMap BRAVO liquid handling system. Cartridges were sequentially conditioned with 100% acetonitrile (ACN) and then water 0.1% FA, samples were then loaded, washed with 0.1% FA, and peptides eluted with 60% ACN, 0.1% FA. Finally, the organic solvent was removed in a SpeedVac concentrator.

### Global Proteomics LC-MS Analysis

Prior to LC-MS/MS analysis, dried peptides were reconstituted with 2% ACN, 0.1% FA and concentration was determined using Stunner UV/Vis plate reader (Unchained Labs). Samples were then brought to the same concentration and analyzed by LC-MS/MS using a nanoElute system (Bruker Daltonics) coupled to a TimsTOF Pro2 mass spectrometer (Bruker Daltonics) via a CaptiveSpray nano-electrospray source (Bruker Daltonics). The source parameters were: 1700 V of capillary voltage, 3.0 L/min of dry gas, and 200°C of dry temperature. Peptides were separated with an analytical reversed-phase C18 Aurora column (25 cm × 75µm i.d., 1.6 µm particles; IonOpticks) at a constant flow rate of 400 nL/min over a 20 minute non-linear gradient from 2 to 34% of mobile phase B (mobile phase A= FA 0.1%; mobile phase B=99.9% ACN: 0.1% FA). Temperature of the analytical column was kept at 50°C. All MS data was acquired in diaPASEF mode with one MS1 full scan was followed by 8 diaPASEF scans with 2 variable widths mobility windows each, making a total of 16 windows that were optimized for sensitivity and reproducibility using py_diAID^88^. The method covers the ion mobility range (1/K0) from 0.69 to 1.43 V s/cm2, and m/z range from 312.9 to 1490.3 Da. The ramp times were set to 100 ms, giving a total cycle time of 0.95 seconds. The collision energy was increased as a function of ion mobility, following a straight line from 20 eV for 1/K0 of 0.6 V s/cm2 to 59 eV for 1/K0 of 1.6 V s/cm2. The TIMS ion mobility dimension was calibrated using three ions of Tuning mix ES-TOF CCS compendium (ESI) (m/z [Th], 1/K0 [Th]: 622.0290, 0.9915; 922.0098, 1.1986; 1221.9906, 1.3934).

### Global proteomics data processing and analysis

All raw files were processed with DIA-NN software^86^ version 1.8.0 using library-free mode against the reviewed Uniprot/SwissProt human proteome concatenated with a MaxQuant contaminant database, giving a total of 20,565 entries. In brief, mass accuracies were set to 15 ppm, and scan window was set to 0 (automatic). MBR was enabled for the analysis and “Robust LC (high precision)” quantification was selected. No normalization was applied at this level. For the remaining settings, the DIA-NN default was used. The report files from DIA-NN were used for statistical analysis carried out utilizing custom scripts in R Statistical Software^77^ and limma^84^ and MSstats^89^ software packages. In brief, all identifications with Global.PG.Q.Value > 0.01 & Global.Q.Value > 0.01 were removed. Feature intensities were log2 transformed and loess-normalized across all samples to account for systematic errors. Testing for differential abundance was performed using MSstats based on a linear mixed-effects model. Prior to statistical test, peptides shared across protein groups were removed. Imputation of missing values of the data was avoided prior to statistical test; therefore, a pseudo log2FC, adj.pvalue and pvalue only was calculated for proteins completely missing in one condition. To calculate the pseudo log2FC, peptide intensities of the one-condition missing protein were summed across all replicates in the group where it was detected, then divided the total sum intensity by 3.3. The imputed pvalue and adj.pvalue was calculated by dividing 0.05 or 0.1, respectively, by the number of replicates that the given protein was confidently identified (in the condition where it is detected) multiplied by the number of features (i.e., precursor ions) quantified. Therefore, the imputed log2FC gives an estimate of the protein abundance in the condition where it was detected, while the imputed pvalue or adj.pvalue reports the confidence in consistency of protein detection for one of the conditions being statistically tested.

### GEMIN3 degradation assay

The HiBiT-GEMIN3 cell line was generated by CRISPR editing and clonal isolation, followed by genomic PCR, Sanger sequencing, and HiBiT assay for validation. Briefly, 0.6 × 10^6^ HEK293T cells were transfected in a 12-well plate using Lipofectamine 3000 and 3 plasmids: CRISPR enzyme with GEMIN3-targeting sgRNA (TGACGGCGCGGCTACCATGG), HDR template, and i53 HDR enhancer^90^. After 6 h, cells were passaged 1:8 and 1 µM M3814 was added to the media as an HDR enhancer. After 24 h, cells were placed under 2 µg/mL puromycin selection for the next two days. Cells were subsequently expanded, and clones were isolated by limiting dilution. For the GEMIN3 degradation assay, validated HiBiT-GEMIN3 HEK293T cells were dispensed in 384-well format with 2,000 cells per well and cultured overnight. Compounds were dosed by acoustic dispensing (Labcyte Echo 555) of 10mM DMSO stocks, and HiBiT-tagged proteins were detected by addition of Nano-Glo HiBiT Lytic Reagent (Promega #N3040) according to the manufacturer specifications and measuring luminescence on a multimode plate reader (EnVision Xcite).

### NanoBiT ternary complex assay

VHL, GEMIN3, and other DEAD-box helicase genes were synthesized, subcloned using SgfI/PmeI restriction sites into NanoBiT PPI Flexi vectors (Promega #N2015) and verified by Sanger sequencing. HEK293T cells were plated at 10k cells per well in 96-well plates in OptiMEM with 5% FBS. After cells adhered to the plates, transfection mixtures were prepared in bulk and each well was individually transfected using 0.3 μL Fugene 6 (Promega #E2691), 50 ng LgBiT-tagged VHL, and 50 ng SmBiT-tagged target in 5 μL OptiMEM. After 24 hour transfection, a 30-minute pre-treatment with 1 μM bortezomib was used to inhibit the proteasome. For competition experiments, the VHL ligand VH298 was also included during the pre-treatment at a concentration of 10 μM. For single-dose experiments, cells were treated with dGEM at a final concentration of 10 μM for 2 hours. For dose-response experiments, dGEM3 was serially diluted in DMSO before preparing a working solution in media that was added to cells. After 2 hour incubation, plates were brought to room temperature and 25 μL of freshly prepared Nano-Glo Live Cell Reagent (Promega #N2012) was added to each well. Luminescence was measured immediately after mixing using an EnVision XCite multimode plate reader (PerkinElmer #2105-010).

### HTRF ternary complex assay

For the HTRF ternary complex assay, a reaction mixture was prepared with 10 nM biotinylated VHL/EloB/EloC complex, 40 nM 6HIS-tagged GEMIN3_aa41-268_ or BRD4_BD2_, and mAb Anti-6HIS-d2 (Revvity #61HISDLF) and Streptavidin-Terbium cryptate (Revvity #610SATLF) at concentrations as specified by the manufacturer. This mixture was dispensed in 384-well format, and compounds were added to 20 μL reactions by acoustic dispensing of 10 mM DMSO stocks. After incubating 1 h, the acceptor and donor emissions were measured using a multimode plate reader (EnVision Xcite), and the background-normalized 665/620 HTRF ratio was calculated. For competition experiments, VH298 was dispensed in dose-response prior to addition of dGEM3 or MZ1.

### Protein purifications

Ubiquitin (#U-100H), UBA1 (#E-305), neddylated CUL2/RBX1 (#E3-421), and VHL/EloB/EloC (#E3-600) were obtained commercially from R&D Systems. The cDNAs encoding 6HIS-tagged UBE2R2, UBE2L3, SUMO-ARIH1, and HiBiT-GEMIN3_aa41-268_ were cloned into the pET28a vector. A cDNA encoding 6HIS-HiBiT-HA-BRD4_BD2_ was cloned into the pMAL-c4x vector. A cDNA encoding GST-6HIS-TEV-TUBE with two additional cysteines was cloned into the pET41 vector. Recombinant proteins were expressed in *E. coli* BL21 (DE3) in autoinduction medium at 18 °C for 16 h. Proteins were first purified using Ni-NTA resin (Cytiva). The aggregates were subsequently removed by gel-filtration column chromatography using Hi-Load Superdex200 16/60 (Cytiva). The N-terminal 6HIS-SUMO tag of ARIH1, MBP-6HIS tag of BRD4_BD2_, and GST-fusion tag were removed by ULP1 and TEV protease digestions, respectively. ARIH1, HiBiT-HA-BRD4, and 6HIS-HiBiT-GEMIN3_aa41-268_ were further purified with anion exchange column chromatography using Capto HiRes Q 10/100 (Cytiva). The TUBE protein was thiol-conjugated with TAMRA (TAMRA Maleimide, BioActs KWM1057) at two cysteine residues and purified over by gel-filtration column chromatography using Hi-Load Superdex200 16/60 (Cytiva). All recombinant proteins were buffer exchanged into storage buffer (25 mM HEPES, pH 7.5, 150 mM NaCl, 10% Glycerol, 1 mM TCEP) using centrifugal concentrators (Amicon) and aliquoted prior to flash-freezing with liquid nitrogen for storage.

### tBRET ubiquitination assay

For the tBRET ubiquitination assay, a reaction mixture was prepared on ice containing 200 nM VHL/EloB/EloC, 200 nM neddylated CUL2/RBX1, 2.5 μM UBE2R2, 2.5 μM UBE2L3, 200nM ARIH1, 250 nM UBA1, 50 μM ubiquitin, and 2.5 mM ATP. HiBiT-GEMIN3_aa41-268_ or HiBiT-HA-BRD4_BD2_ was added to this mixture at a final concentration of 1 μM, mixed, and dispensed in 384-well format. Compounds were added by acoustic dispensing of 10 mM DMSO stocks (Echo 555) and incubated at room temperature for 30 minutes. For plate-based detection of ubiquitination, a tBRET detection mixture was prepared with 0.625 μM TUBE-TAMRA protein in a dilution of 1:200 LgBiT protein and 1:100 Nano-Glo Luciferase Assay Substrate (Promega #N2410). For each replicate, 2.5 μL of the reaction mixture was diluted 1:20 into tBRET detection mixture, shaken at room temperature for 5 minutes, and read on a multimode plate reader. For western blot detection of HiBiT, the remaining reaction mixture was diluted 1:1 in SDS sample buffer, run on a 4-20% SDS-PAGE gel, and imaged using the NanoGlo HiBiT Blotting system according to manufacturer specifications (Promega #N2410).

### SPR

SPR experiments were performed on a Biacore 8K or T200 instrument (Cytiva) using previously established protocols^76^. Immobilization of His-tagged VHL/EloB/EloC (VBC) or GEMIN3_41-268_ was carried out at 25°C using Series S NTA chip, where protein was first captured via His/NTA affinity followed by amine-coupling using NHS/EDC, and finally deactivation of the unfunctionalized carboxy groups using 1 M ethanolamine. The running buffer during immobilization was 20 mM HEPES, 200 mM NaCl with 0.005% P20. High-density (4000-5000 RU), medium-density (1000-2000 RU), and low density (500-1000 RU) GEMIN3- or VBC-functionalized surfaces were created, followed by equilibration in the running buffer for 3 h. For ternary complex analysis, compounds (10 mM diluted to 100μM in 100% DMSO) were initially prepared at 400 nM in a running buffer (200mM NaCl, 20mM HEPES, 1mM PEG-8000, 1mM TCEP, 0.2mg/mL BSA, 0.02% P20). This solution was mixed at 1:1 ratio with a solution of 6 μM untagged VBC in the running buffer. This complex was then serially diluted in the running buffer (6-point two-fold serial dilution). Solutions were injected sequentially in multi-cycle kinetic format without regeneration (contact time 120 s, flow rate 80 μL/min, dissociation time 900 s) using a stabilization period of 60 s on a low-density GEMIN3-functionalized surface. For binary binding analysis, compounds were prepared at 400 uM in DMSO, then diluted to final concentration (25, 50, 100, 200, 400 nM) in running buffer. Compound samples were injected sequentially in multi-cycle kinetic format (contact time [90 s], flow rate [80 μL/min], dissociation time 180 s) using a stabilization period of 60 s on a high-density GEMIN3- or VHL-functionalized surface. For VHL:GEMIN3 native interaction analysis, VBC was diluted in running buffer to different final concentrations (20, 10, 5, 2.5, 1.25, 0.625 uM) and injected in multi-cycle kinetic format (association time 90 s, flow rate 80 μL/min, dissociation time 180 s) using a stabilization period of 60s on a low-density GEMIN3-functionalized surface. Raw sensorgrams were processed by performing reference subtraction, solvent correction, and analysis was done using Biacore Insight Evaluation Software. Kinetic analysis was performed by fitting data to a 1:1 Langmuir interaction model and a steady-state affinity model was used to evaluate equilibrium binding affinity.

## Chemistry Supplement

### Synthesis scheme of compound 1

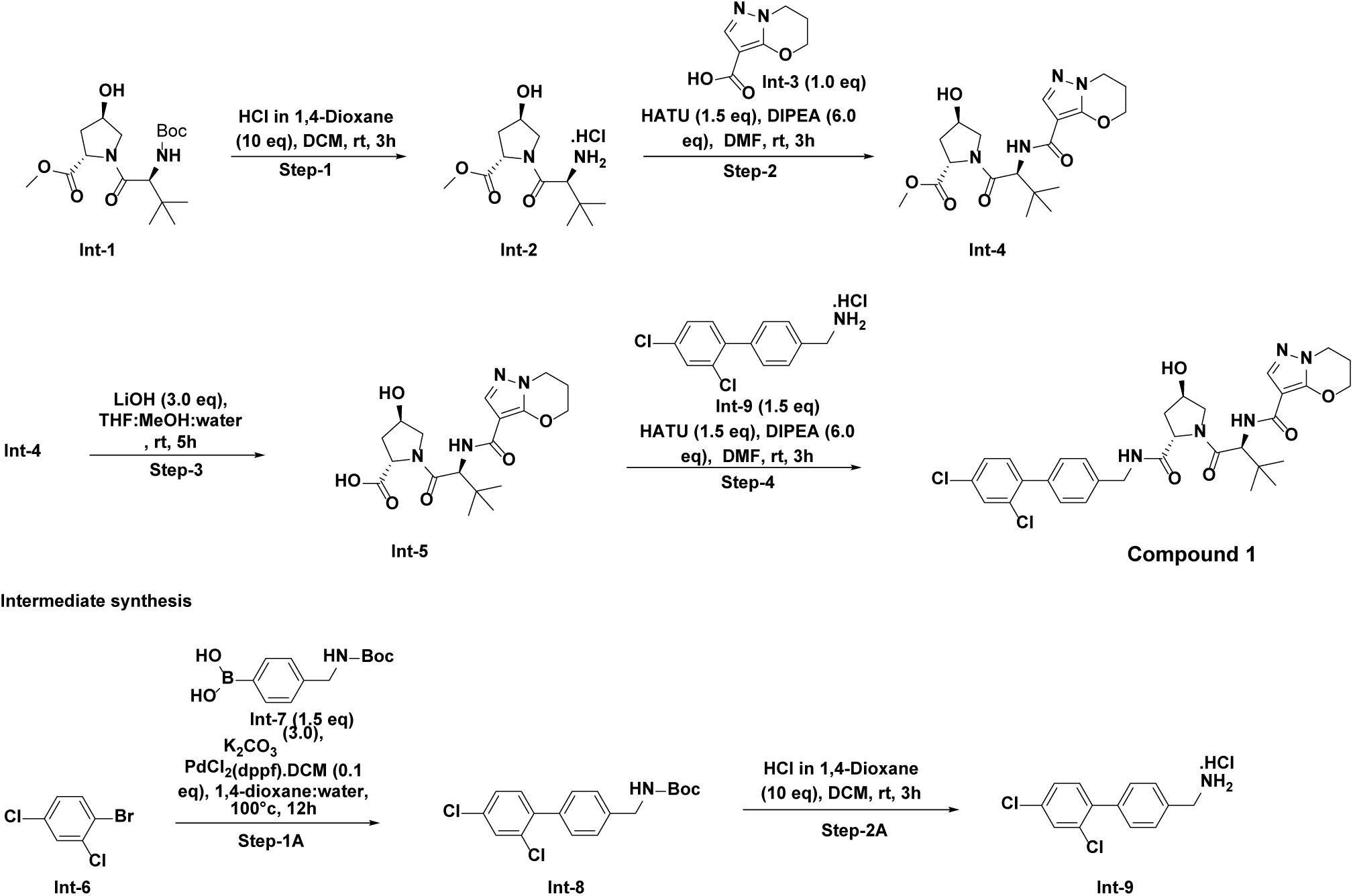

### Experimental procedure of compound 1

Step-1: Synthesis of methyl (2S,4R)-1-((S)-2-amino-3,3-dimethylbutanoyl)-4-hydroxypyrrolidine-2-carboxylate hydrochloride (Int-2)

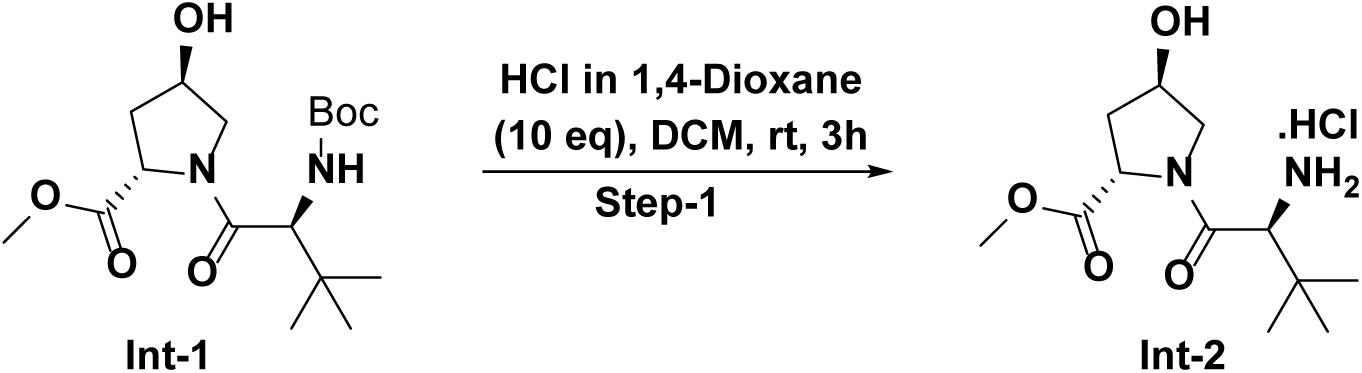

To a 50-mL round-bottomed flask was added methyl (2S,4R)-1-((S)-2-((tert-butoxycarbonyl) amino)-3,3-dimethylbutanoyl)-4-hydroxypyrrolidine-2-carboxylate (600 mg, 1.674 mmol, 1.0 eq) in dichloromethane (20 mL) followed by addition of HCl in 1,4 dioxane (4M) (4185 µL, 16.74 mmol, 10 eq) at 0°C and stirred at room temperature for 5 h. After completion, the reaction mixture was concentrated and the crude, methyl (2S,4R)-1-((S)-2-amino-3,3-dimethylbutanoyl)-4-hydroxypyrrolidine-2-carboxylate hydrochloride (482 mg) was used as such for next step.

MS (ESI, Positive & negative ion) *m/z*: 259.1 (M+1)

Step-2: Synthesis of methyl (2S,4R)-1-((S)-2-(6,7-dihydro-5H-pyrazolo[5,1-b] [1,3] oxazine-3-carboxamido)-3,3-dimethylbutanoyl)-4-hydroxypyrrolidine-2-carboxylate (Int-4)

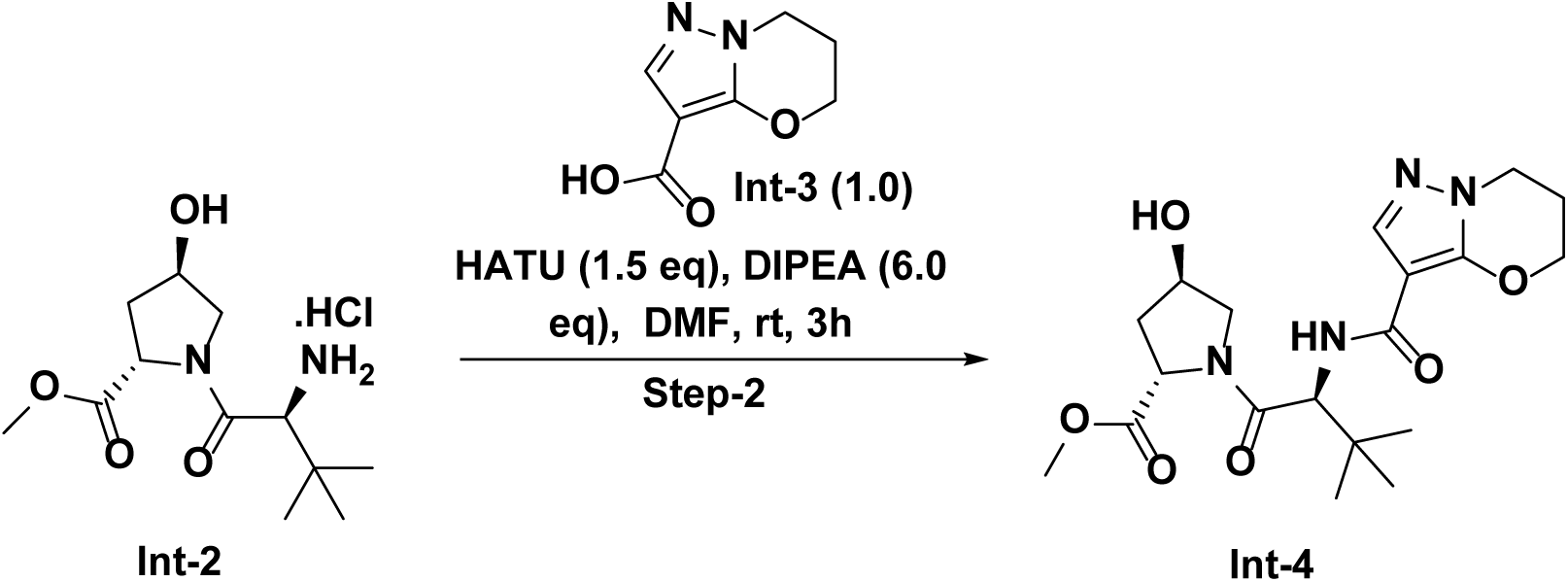

To a stirred solution of 6,7-dihydro-5H-pyrazolo[5,1-b] [1,3] oxazine-3-carboxylic acid (250 mg, 1.487 mmol, 1.0 eq) in N, N-dimethylformamide (10 mL) was added methyl (2S,4R)-1-((S)-2-amino-3,3-dimethylbutanoyl)-4-hydroxypyrrolidine-2-carboxylate hydrochloride (482 mg, 1.635 mmol, 1.1 eq), DIPEA (1298 µL, 7.43 mmol, 6.0 eq). and HATU (848 mg, 2.230 mmol, 1.5 eq) at 0°C and stirred at room temperature for 2 h. After completion, the reaction mixture was concentrated and the crude material was absorbed onto a plug of silica gel and purified by flash column chromatography through a Redi-Sep pre-packed silica gel column (12 g), eluting with a gradient of 3% to 5% MeOH in CH_2_Cl_2_, to provide methyl (2S,4R)-1-((S)-2-(6,7-dihydro-5H-pyrazolo[5,1-b] [1,3] oxazine-3-carboxamido)-3,3-dimethylbutanoyl)-4-hydroxypyrrolidine-2-carboxylate (500 mg, 1.224 mmol, 82 % yield) as an off-white solid.

MS (ESI, Positive & negative ion) *m/z*: 409.2 (M+1)

Step-3: Synthesis of (2S,4R)-1-((S)-2-(6,7-dihydro-5H-pyrazolo[5,1-b] [1,3] oxazine-3-carboxamido)-3,3-dimethylbutanoyl)-4-hydroxypyrrolidine-2-carboxylic acid (Int-5)

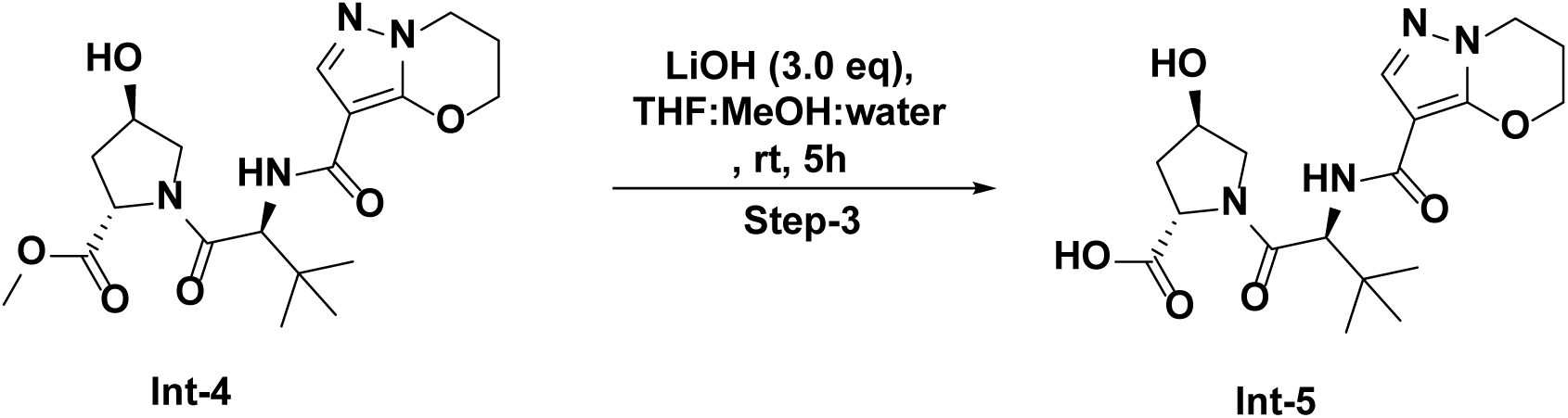

To a stirred soultion methyl (2S,4R)-1-((S)-2-(6,7-dihydro-5H-pyrazolo[5,1-b] [1,3] oxazine-3-carboxamido)-3,3-dimethylbutanoyl)-4-hydroxypyrrolidine-2-carboxylate (600 mg, 1.469 mmol, 1.0 eq) in tetrahydrofuran (14 mL) and methanol (2 mL), was added a solution of lithium hydroxide hydrate (123 mg, 2.94 mmol, 2.0 eq) in water (4 mL) at 0°C and stirred at 10-15°C for 5 h. After completion, the reaction mixture was concentrated and diluted with water (10 mL) and extracted with ethyl acetate (2 × 10 mL). The aqueous layer pH was adjusted to 4-5 using 1.5 M HCl solution and extracted with 10% methanol in DCM (3 × 30 mL), dried over Na_2_SO_4_. The solution was filtered and concentrated in vacuum to give the crude (2S,4R)-1-((S)-2-(6,7-dihydro-5H-pyrazolo[5,1-b] [1,3] oxazine-3-carboxamido)-3,3-dimethylbutanoyl)-4-hydroxypyrrolidine-2-carboxylic acid (435 mg, 1.102 mmol, 75 % yield) as an off-white solid.

MS (ESI, Positive & negative ion) *m/z*: 394.2 (M+1)

^1^H NMR (DMSO-*d*_6_, 400 MHz): δ (ppm) 12.46 (s, 1H), 7.60 (s, 1H), 6.81 (d, *J*=9.4 Hz, 1H), 5.18 (d, *J*=3.7 Hz, 1H), 4.67 (d, *J*=9.4 Hz, 1H), 4.52 (t, *J*=5.6 Hz, 2H), 4.34 (d, *J*=4.3 Hz, 1H), 4.26 (dd, *J*=9.2, 7.7 Hz, 1H), 4.10 (t, *J*=6.1 Hz, 2H), 3.72 – 3.61 (m, 2H), 2.24 (dd, *J*=7.1, 4.4 Hz, 2H), 2.11 (t, *J*=10.5 Hz, 1H), 1.96 – 1.85 (m, 2H), 0.97 (s, 9H).

Step-4: Synthesis of N-((S)-1-((2S,4R)-2-(((2’,4’-dichloro-[1,1’-biphenyl]-4-yl) methyl) carbamoyl)-4-hydroxypyrrolidin-1-yl)-3,3-dimethyl-1-oxobutan-2-yl)-6,7-dihydro-5H-pyrazolo[5,1-b] [1,3] oxazine-3-carboxamide (Compound 1)

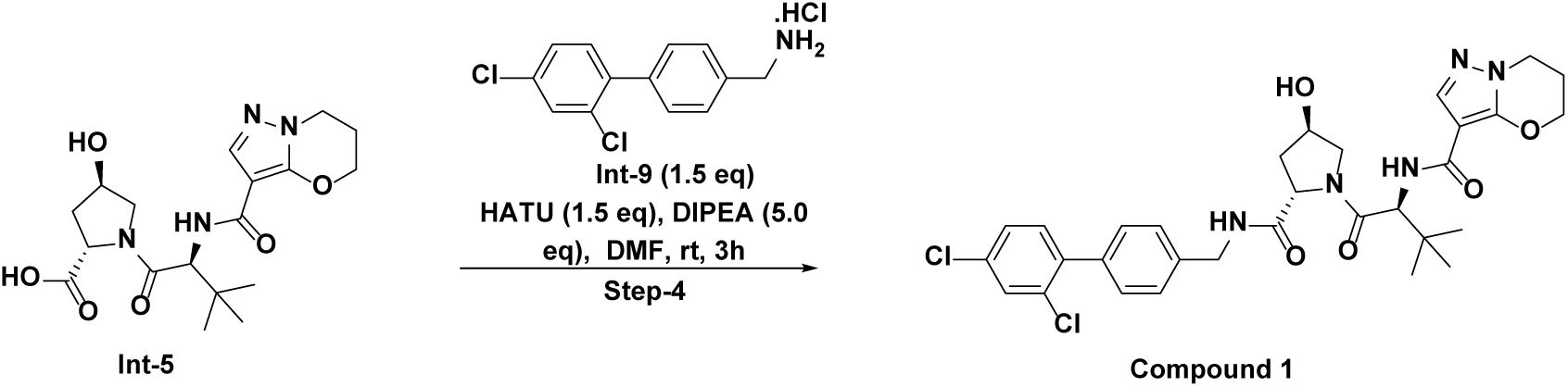

To a stirred solution of (2S,4R)-1-((S)-2-(6,7-dihydro-5H-pyrazolo[5,1-b] [1,3] oxazine-3-carboxamido)-3,3-dimethylbutanoyl)-4-hydroxypyrrolidine-2-carboxylic acid (200 mg, 0.507 mmol, 1.0 eq) and (2’,4’-dichloro-[1,1’-biphenyl]-4-yl) methanamine hydrochloride (220 mg, 0.761 mmol, 1.5 eq) in N, N-dimethylformamide (10 mL), was added DIPEA (443 µL, 2.54 mmol, 5.0 eq) followed by HATU (289 mg, 0.761 mmol, 1.5 eq) at 0°C. and stirred at room temperature for 2 h. TLC (10% methanol in DCM, Rf-0.5) showed completion of the starting material and non-polar spot detected. The reaction mixture was concentrated and the crude material was absorbed onto a plug of silica gel and purified by reverse phase chromatography through a C18 Redi-Sep pre-packed column (40 g), eluting with a gradient of 55% to 60% acetonitrile in water, to provide N-((S)-1-((2S,4R)-2-(((2’,4’-dichloro-[1,1’-biphenyl]-4-yl) methyl) carbamoyl)-4-hydroxypyrrolidin-1-yl)-3,3-dimethyl-1-oxobutan-2-yl)-6,7-dihydro-5H-pyrazolo[5,1-b] [1,3] oxazine-3-carboxamide (191 mg, 3.038 mmol, 60 % yield) as a white solid.

MS (ESI, Positive & negative ion) *m/z*: 628.2 (M+1)

^1^H NMR (Methanol-*d*_4_, 400 MHz): δ (ppm) 7.71 (s, 1H), 7.57 (d, *J*=2.0 Hz, 1H), 7.47 – 7.42 (m, 2H), 7.41 – 7.34 (m, 4H), 7.14 (d, *J*=9.2 Hz, 1H), 4.82 (d, *J*=9.2 Hz, 1H), 4.64 – 4.52 (m, 5H), 4.47 – 4.39 (m, 1H), 4.19 (t, *J*=6.1 Hz, 2H), 3.97 (d, *J*=11.1 Hz, 1H), 3.85 (dd, *J*=11.0, 3.9 Hz, 1H), 2.35 (td, *J*=5.5, 3.5 Hz, 2H), 2.26 (dd, *J*=13.2, 7.6 Hz, 1H), 2.12 (ddd, *J*=13.3, 9.2, 4.5 Hz, 1H), 2.03 (s, 1H), 1.09 (s, 9H).

^13^C NMR (151 MHz, DMSO-d_6_) d ppm 21.40 (s, 1 C) 23.05 (s, 1 C) 26.69 (s,3 C) 36.21 (s, 1C) 38.41 (s, 1 C) 42.20 (s, 1 C) 44.22 (s, 1 C) 55.92 (s, 1 C) 57.02 (s, 1 C) 59.18 (s, 1 C) 67.74 (s,1 C) 69.34 (s, 1 C) 98.26 (s, 1 C) 127.37 (s, 2 C) 127.66 (s, 1 C) 128.14 (s, 1 C) 129.43 (s, 2 C) 129.58 (s, 1 C) 129.72 (s, 1 C) 132.82 (s, 1 C) 133.16(s,1 C) 133.25 (s, 1 C) 136.37 (s, 1 C) 138.94 (s, 1 C) 139.12 (s, 1 C) 139.91 (s, 1 C) 149.88 (s, 1 C) 160.80 (s, 1 C) 170.08 (s, 1 C) 172.20 (s, 1 C)

### Experimental procedure of Int-9

Step-1A: Synthesis of tert-butyl ((2’,4’-dichloro-[1,1’-biphenyl]-4-yl) methyl) carbamate (Int-8)

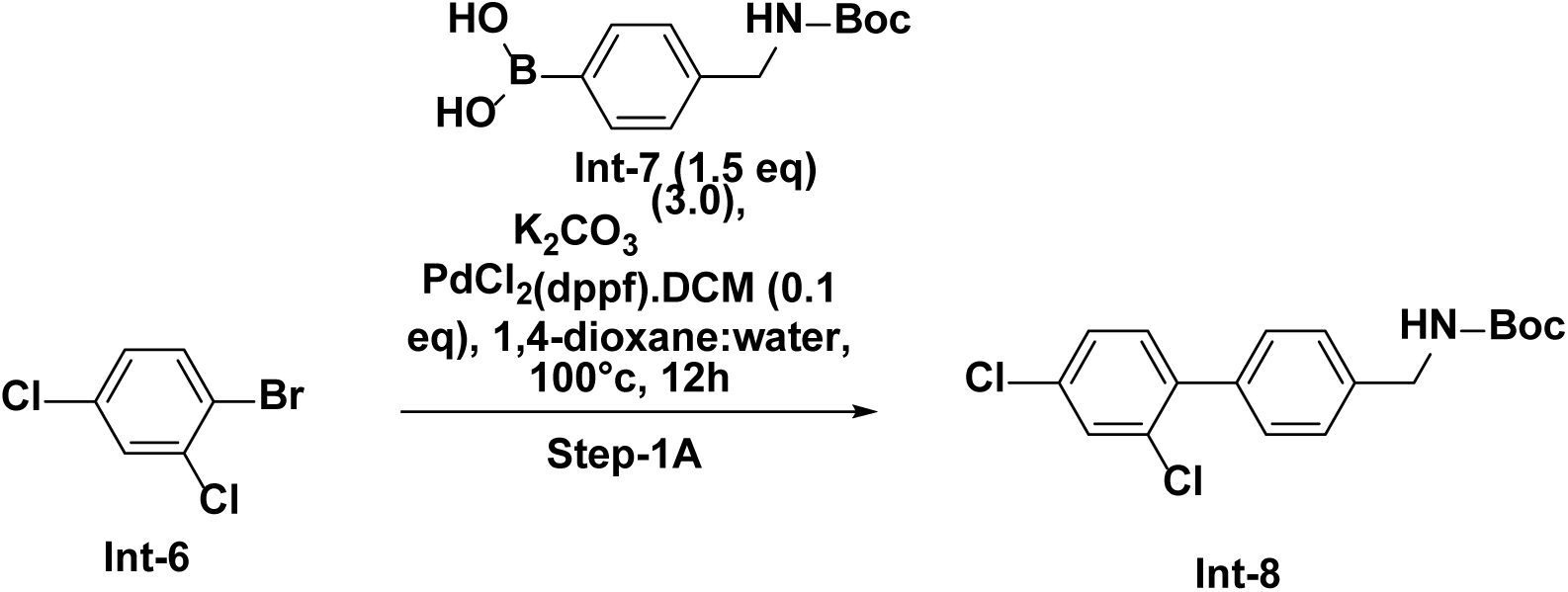

To a 20 mL Wheaton vial was added 1-bromo-2,4-dichlorobenzene (250 mg, 1.107 mmol, 1.0) and (4-(((tert-butoxycarbonyl) amino) methyl) phenyl) boronic acid (417 mg, 1.660 mmol, 1.5 eq), K_2_CO_3_ (459 mg, 3.32 mmol, 3.0 eq),1,4-dioxane (8.0 mL) and water (2.0 mL). After degassing, PdCl_2_(dppf)-CH_2_Cl_2_ adduct (90 mg, 0.111 mmol, 0.1 eq) was added and the reaction mixture was stirred at 100 °C for 12 h. TLC (30% ethyl acetate in hexane, Rf-0.5) showed completion of the starting material and one major nonpolar spot detected. The reaction mixture was filtered through celite and concentrated, absorbed onto a plug of silica gel and purified by column chromatography through a Redi-Sep pre-packed silica gel column (12 g), eluting with a gradient of 5% to 10% EtOAc in hexane, to provide tert-butyl ((2’,4’-dichloro-[1,1’-biphenyl]-4-yl) methyl) carbamate (320 mg, 0.908 mmol, 82 % yield) as an off-white solid.

MS (ESI, Positive & negative ion) *m/z*: 386.1 (M+1)

^1^H NMR (400 MHz, DMSO-*d*_6_) δ 7.73 (d, *J* = 2.1 Hz, 1H), 7.53 – 7.50 (m, 1H), 7.44 (t, *J* = 8.8 Hz, 2H), 7.38 (d, *J* = 8.2 Hz, 2H), 7.34 (s, 2H), 4.19 (d, *J* = 6.3 Hz, 2H), 1.41 (s, 9H).

Step-2A: Synthesis of (2’,4’-dichloro-[1,1’-biphenyl]-4-yl) methanamine hydrochloride (Int-9)

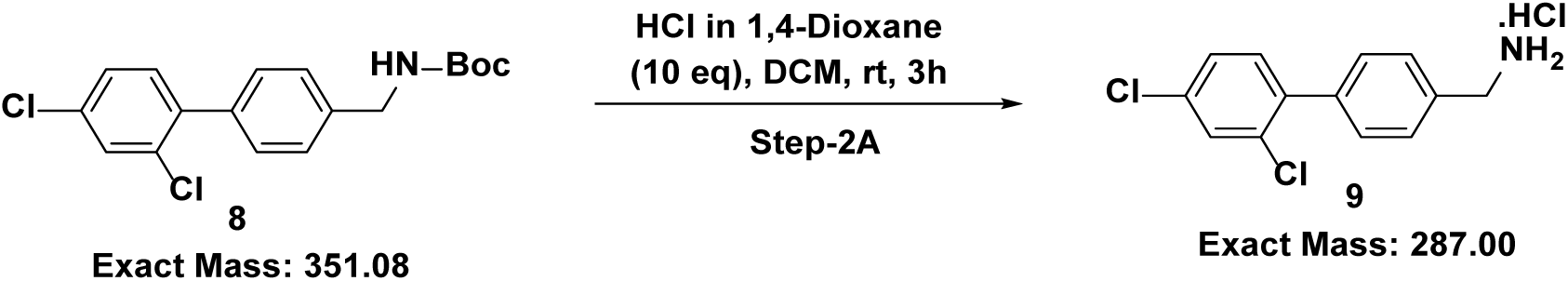

To a stirred solution of tert-butyl ((2’,4’-dichloro-[1,1’-biphenyl]-4-yl) methyl) carbamate (250 mg, 0.710 mmol, 1.0 eq) in dichloromethane (10 mL), was added HCl in 1,4-dioxane (4M) (1774 µL, 7.10 mmol, 10 eq) at 0°C and the reaction mixture was stirred at room temperature for 5 h. After completion, the reaction mixture was concentrated and the crude (2’,4’-dichloro-[1,1’-biphenyl]-4-yl) methanamine hydrochloride (215 mg) used as such for next step.

MS (ESI, Positive & negative ion) *m/z*: 253.2 (M+1)

### Synthesis scheme of compound 3

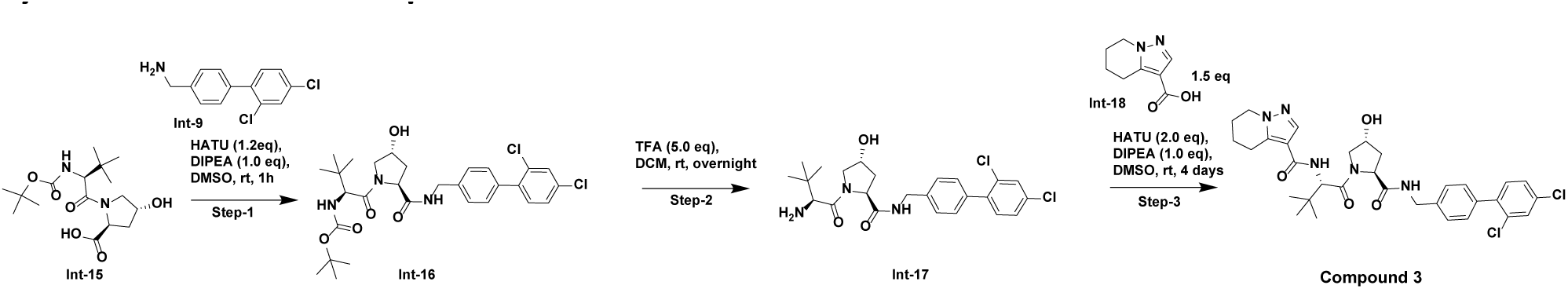

### Experimental procedure of compound 3

Step-1: Synthesis of tert-butyl ((S)-1-((2S,4R)-2-(((2’,4’-dichloro-[1,1’-biphenyl]-4-yl)methyl)carbamoyl)-4-hydroxypyrrolidin-1-yl)-3,3-dimethyl-1-oxobutan-2-yl)carbamate (Int-16)

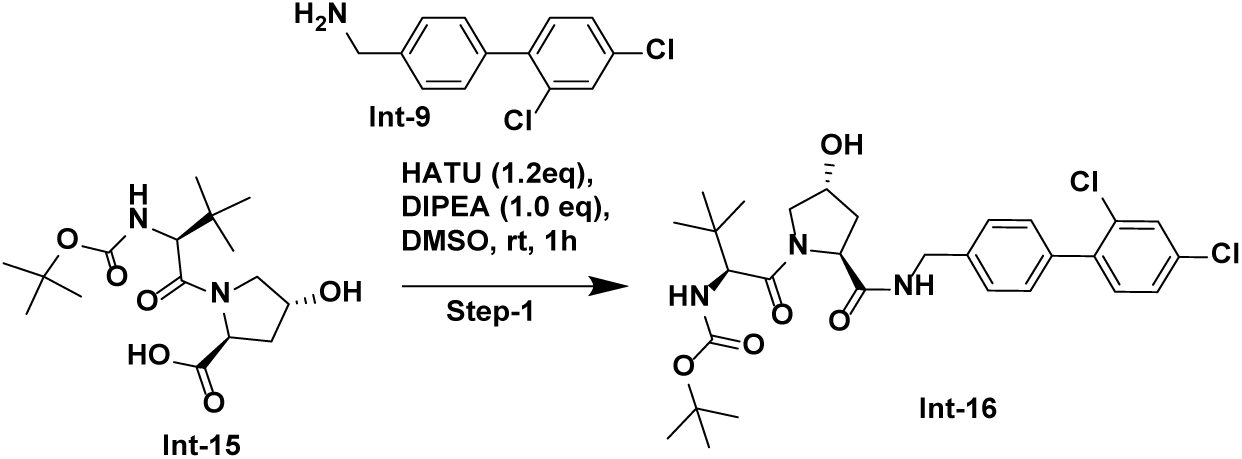

To a 4mL dram vial was added l-proline, n-[(1,1-dimethylethoxy)carbonyl]-3-methyl-l-valyl-4-hydroxy-, (4r)-(72 mg, 0.209 mmol) and HATU (95 mg, 0.251 mmol) in dimethyl sulfoxide (.5 mL) followed by (2’,4’-dichloro-[1,1’-biphenyl]-4-yl)methanamine (52.7 mg, 0.209 mmol) and 1,1’-dimethyltriethylamine (27.0 mg, 0.037 mL, 0.209 mmol) and stirred at room temperature for 1 hour. The reaction mixture was purified by prep-HPLC on a Phenomenex Luna column, 5 micron, C8(2), 100 Å, 150 × 21.2 mm eluting at 20 mL/min with a gradient of 10-100% acetonitrile (0.1% TFA) in acetonitrile (0.1% TFA). The combined fractions containing the product were frozen at -78 °C, and lyophilized. tert-butyl ((S)-1-((2S,4R)-2-(((2’,4’-dichloro-[1,1’-biphenyl]-4-yl)methyl)carbamoyl)-4-hydroxypyrrolidin-1-yl)-3,3-dimethyl-1-oxobutan-2-yl)carbamate (56.7 mg, 0.098 mmol, Yield: 46.9 %, Purity: 99%) was obtained as a solid.

Step-2: Synthesis of (2S,4R)-1-((S)-2-amino-3,3-dimethylbutanoyl)-N-((2’,4’-dichloro-[1,1’-biphenyl]-4-yl)methyl)-4-hydroxypyrrolidine-2-carboxamide (Int-17)

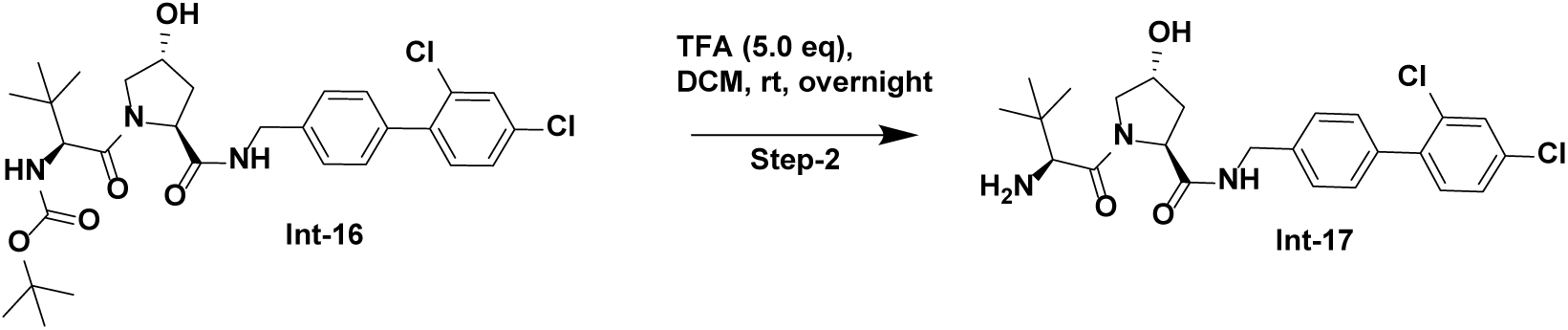

To a 4mL dram vial was added tert-butyl ((S)-1-((2S,4R)-2-(((2’,4’-dichloro-[1,1’-biphenyl]-4-yl)methyl)carbamoyl)-4-hydroxypyrrolidin-1-yl)-3,3-dimethyl-1-oxobutan-2-yl)carbamate (57 mg, 0.099 mmol) in dichloromethane (.5mL) followed by trifluoroacetic acid (56.2 mg, 0.056 mL, 0.493 mmol) and stirred at room temperature for 1 hour. After completion, the reaction mixture was quenched with sodium bicarbonate (5mL). The aqueous layer was extracted in DCM (2 × 10 mL) then dried with Mg_2_SO_4_. The solution was filtered and dried overnight in open air to give the crude (2S,4R)-1-((S)-2-amino-3,3-dimethylbutanoyl)-N-((2’,4’-dichloro-[1,1’-biphenyl]-4-yl)methyl)-4-hydroxypyrrolidine-2-carboxamide (13 mg, 0.027 mmol, 27.6 % yield).

Step-3: Synthesis of N-((S)-1-((2S,4R)-2-(((2’,4’-dichloro-[1,1’-biphenyl]-4-yl)methyl)carbamoyl)-4-hydroxypyrrolidin-1-yl)-3,3-dimethyl-1-oxobutan-2-yl)-4,5,6,7-tetrahydropyrazolo[1,5-a]pyridine-3-carboxamide (Compound 3)

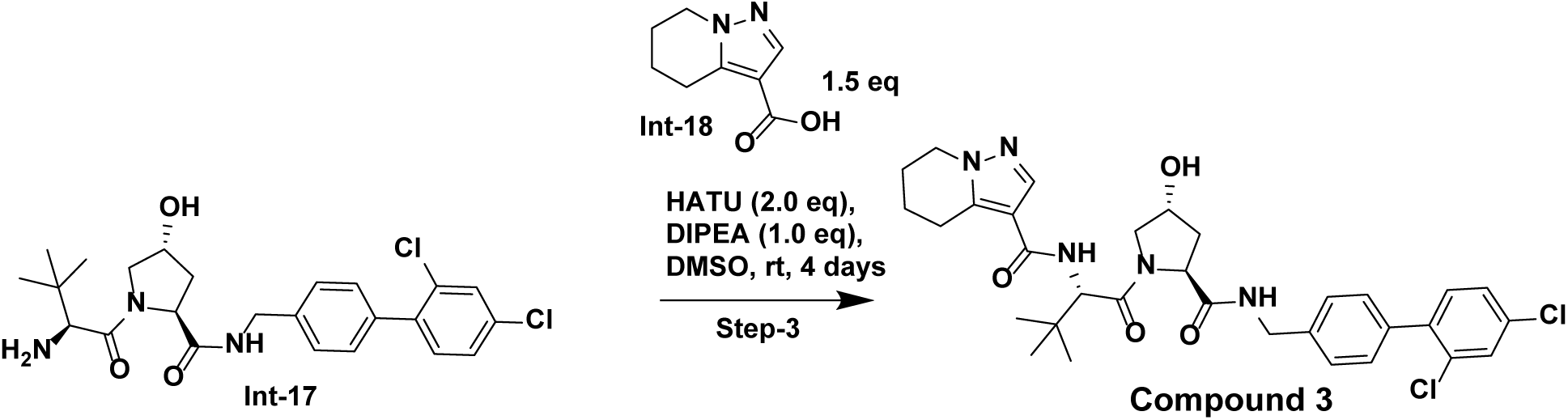

To a 4mL dram vial was added (2S,4R)-1-((S)-2-amino-3,3-dimethylbutanoyl)-N-((2’,4’-dichloro-[1,1’-biphenyl]-4-yl)methyl)-4-hydroxypyrrolidine-2-carboxamide (13 mg, 0.027 mmol), 4,5,6,7-tetrahydropyrazolo[1,5-a]pyridine-3-carboxylic acid (6.77 mg, 0.041 mmol), 1,1’-dimethyltriethylamine (3.51 mg, 4.75 µL, 0.027 mmol), HATU (20.66 mg, 0.054 mmol in dimethyl sulfoxide and stirred at room temperature for 4 days. The reaction mixture was purified by prep-HPLC on a Phenomenex Luna column, 5 micron, C8(2), 100 Å, 150 × 21.2 mm eluting at 20 mL/min with a gradient of 10-100% acetonitrile (0.1% FA) in acetonitrile (0.1% FA). The combined fractions containing the product were frozen at -78 °C, and lyophilized. N-((S)-1-((2S,4R)-2-(((2’,4’-dichloro-[1,1’-biphenyl]-4-yl)methyl)carbamoyl)-4-hydroxypyrrolidin-1-yl)-3,3-dimethyl-1-oxobutan-2-yl)-4,5,6,7-tetrahydropyrazolo[1,5-a]pyridine-3-carboxamide (14.5 mg, 0.023 mmol, 85 % yield, 99% purity) was obtained as a white solid.

1H NMR (DMSO-d6, 600 MHz) δ 8.60 (t, 1H, *J*=6.1 Hz), 7.74 (d, 1H, *J*=2.2 Hz), 7.61 (s, 1H), 7.52 (dd, 1H, *J*=2.2, 8.3 Hz), 7.42 (d, 2H, *J*=8.3 Hz), 7.4-7.4 (m, 2H), 7.3-7.4 (m, 2H), 6.83 (d, 1H, *J*=9.3 Hz), 5.15 (d, 1H, *J*=3.5 Hz), 4.69 (d, 1H, *J*=9.5 Hz), 4.5-4.6 (m, 2H), 4.44 (t, 1H, *J*=8.2 Hz), 4.37 (br d, 1H, *J*=5.8 Hz), 4.33 (br d, 1H, *J*=5.8 Hz), 4.10 (t, 2H, *J*=6.1 Hz), 3.6-3.8 (m, 2H), 2.24 (br dd, 2H, *J*=5.4, 10.5 Hz), 2.07 (br dd, 1H, *J*=7.9, 12.5 Hz), 1.9-2.0 (m, 1H), 1.90 (s, 1H), 0.97 (s, 10H)

13C NMR (151 MHz, DMSO-*d*6) δ ppm 21.40 (s, 1 C) 23.05 (s, 1 C) 26.69 (s, 3 C) 36.21 (s, 1 C) 38.41 (s, 1 C) 42.20 (s, 1 C) 44.22 (s, 1 C) 55.92 (s, 1 C) 57.02 (s, 1 C) 59.18 (s, 1 C) 67.74 (s, 1 C) 69.34 (s, 1 C) 98.26 (s, 1 C) 127.37 (s, 2 C) 127.66 (s, 1 C) 128.14 (s, 1 C) 129.43 (s, 2 C) 129.58 (s, 1 C) 129.72 (s, 1 C) 132.82 (s, 1 C) 133.16 (s, 1 C) 133.25 (s, 1 C) 136.37 (s, 1 C) 138.94 (s, 1 C) 139.12 (s, 1 C) 139.91 (s, 1 C) 149.88 (s, 1 C) 160.80 (s, 1 C) 170.08 (s, 1 C) 172.20 (s, 1 C)

### Synthesis scheme of compounds 4-6

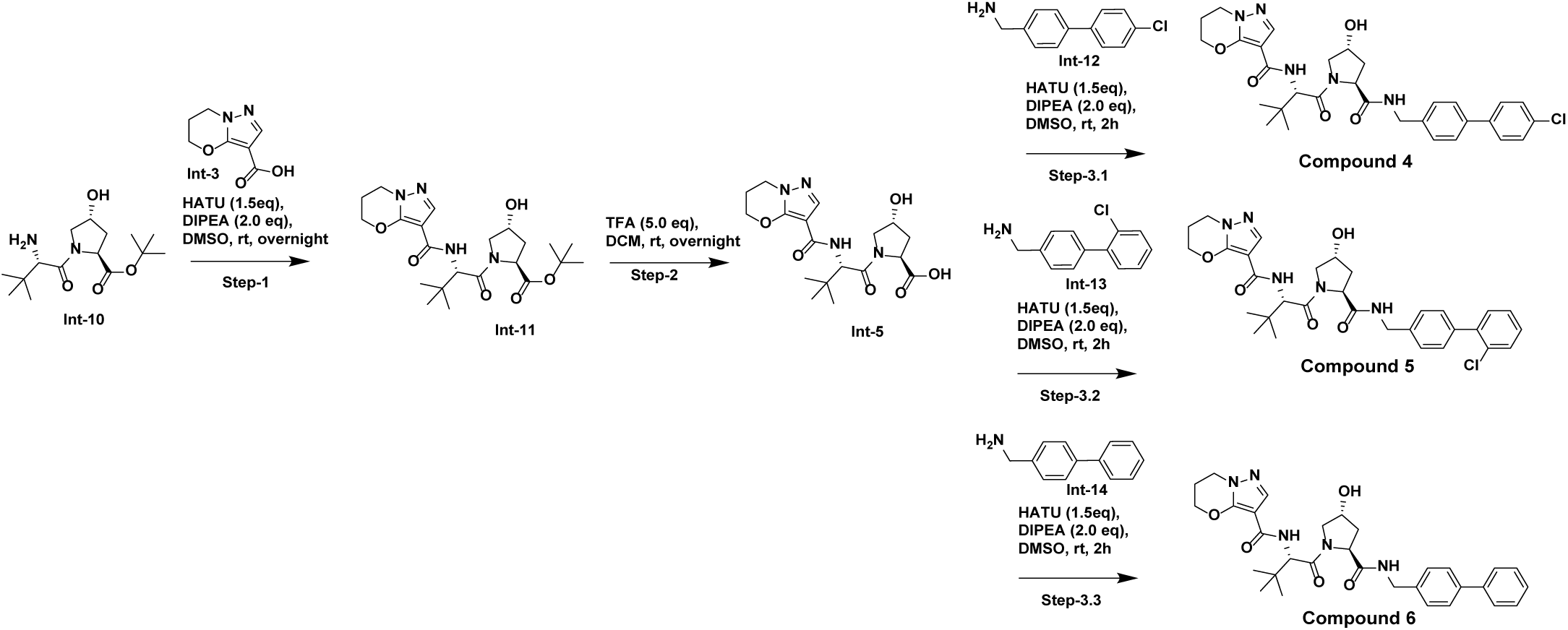

### Experimental procedure of compounds 4-6

Step-1: Synthesis of tert-butyl (2S,4R)-1-((S)-2-(6,7-dihydro-5H-pyrazolo[5,1-b][1,3]oxazine-3-carboxamido)-3,3-dimethylbutanoyl)-4-hydroxypyrrolidine-2-carboxylate (Int-11)

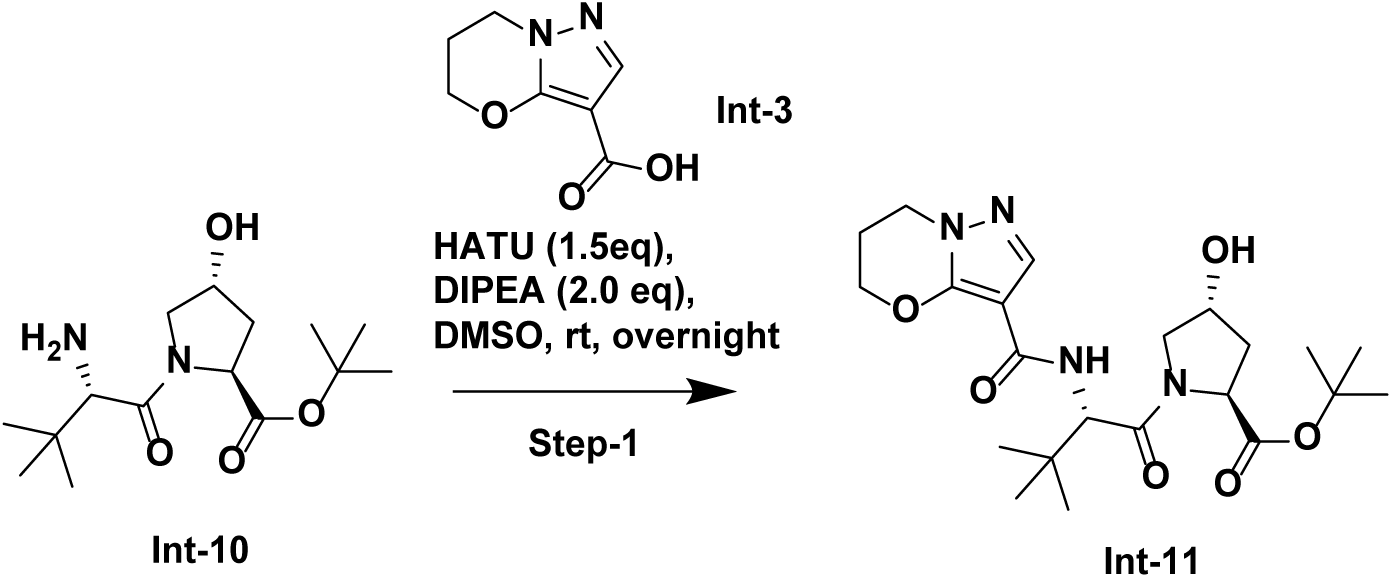

To a 4mL dram vial was added tert-butyl (2S,4R)-1-((S)-2-amino-3,3-dimethylbutanoyl)-4-hydroxypyrrolidine-2-carboxylate (127 mg, 0.423 mmol), 6,7-dihydro-5H-pyrazolo[5,1-b][1,3]oxazine-3-carboxylic acid (89 mg, 0.528 mmol), HATU (241 mg, 0.634 mmol) in dimethyl sulfoxide (2mL), followed by the addition of 1,1’-dimethyltriethylamine (109 mg, 0.148 mL, 0.846 mmol) and stirred at room temperature overnight. The reaction mixture was purified by prep-HPLC on a Phenomenex Luna column, 5 micron, C8(2), 100 Å, 150 × 21.2 mm eluting at 20 mL/min with a gradient of 10-100% acetonitrile (0.1% TFA) in acetonitrile (0.1% TFA). The combined fractions containing the product were frozen at -78 °C, and lyophilized. Tert-butyl (2S,4R)-1-((S)-2-(6,7-dihydro-5H-pyrazolo[5,1-b][1,3]oxazine-3-carboxamido)-3,3-dimethylbutanoyl)-4-hydroxypyrrolidine-2-carboxylate (189 mg, 0.420 mmol, Yield: 99.0 %, Purity: 99 %) was obtained as a solid.

Step-2: Synthesis of (2S,4R)-1-((S)-2-(6,7-dihydro-5H-pyrazolo[5,1-b][1,3]oxazine-3-carboxamido)-3,3-dimethylbutanoyl)-4-hydroxypyrrolidine-2-carboxylic (Int-5):

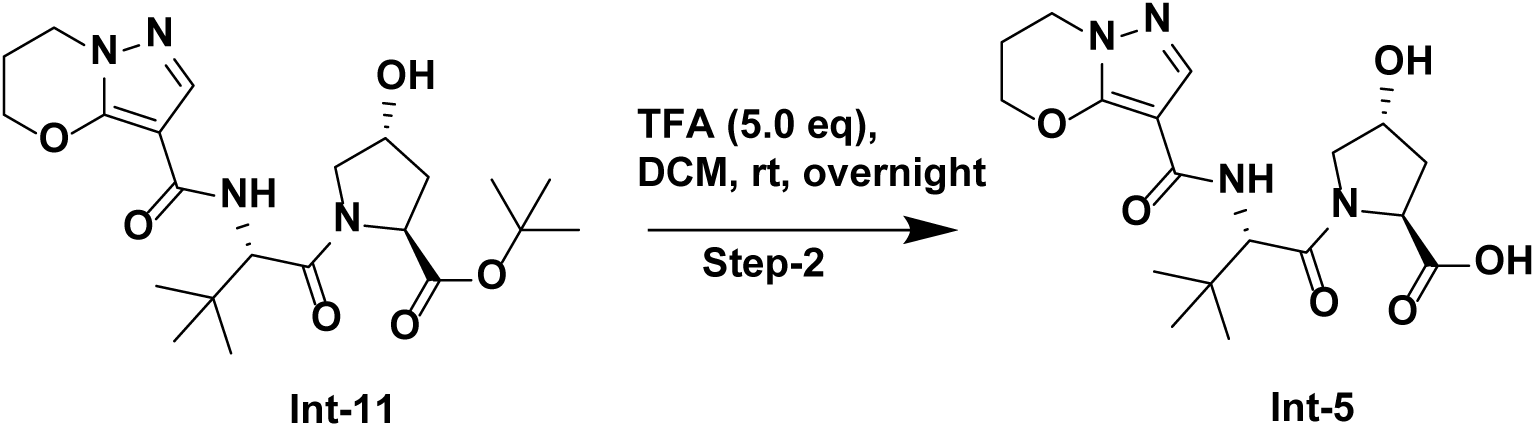

To a 4mL dram vial was added tert-butyl (2S,4R)-1-((S)-2-(6,7-dihydro-5H-pyrazolo[5,1-b][1,3]oxazine-3-carboxamido)-3,3-dimethylbutanoyl)-4-hydroxypyrrolidine-2-carboxylate (189 mg, 0.420 mmol) in dichloromethane (5 mL) followed by trifluoroacetic acid (239 mg, 0.162 mL, 2.098 mmol) and stirred at room temperature overnight. The reaction mixture was concentrated and redissolved in dimethyl sulfoxide then purified by prep-HPLC on a Phenomenex Luna column, 5 micron, C8(2), 100 Å, 150 × 21.2 mm eluting at 20 mL/min with a gradient of 10-100% acetonitrile (0.1% TFA) in acetonitrile (0.1% TFA). The combined fractions containing the product were frozen at -78 °C, and lyophilized. (2S,4R)-1-((S)-2-(6,7-dihydro-5H-pyrazolo[5,1-b][1,3]oxazine-3-carboxamido)-3,3-dimethylbutanoyl)-4-hydroxypyrrolidine-2-carboxylic acid (65 mg, 0.165 mmol, Yield: 39.3 %) obtained.

Step-3-1: Synthesis N-((S)-1-((2S,4R)-2-(((4’-chloro-[1,1’-biphenyl]-4-yl)methyl)carbamoyl)-4-hydroxypyrrolidin-1-yl)-3,3-dimethyl-1-oxobutan-2-yl)-6,7-dihydro-5H-pyrazolo[5,1-b][1,3]oxazine-3-carboxamide (Compound 4)

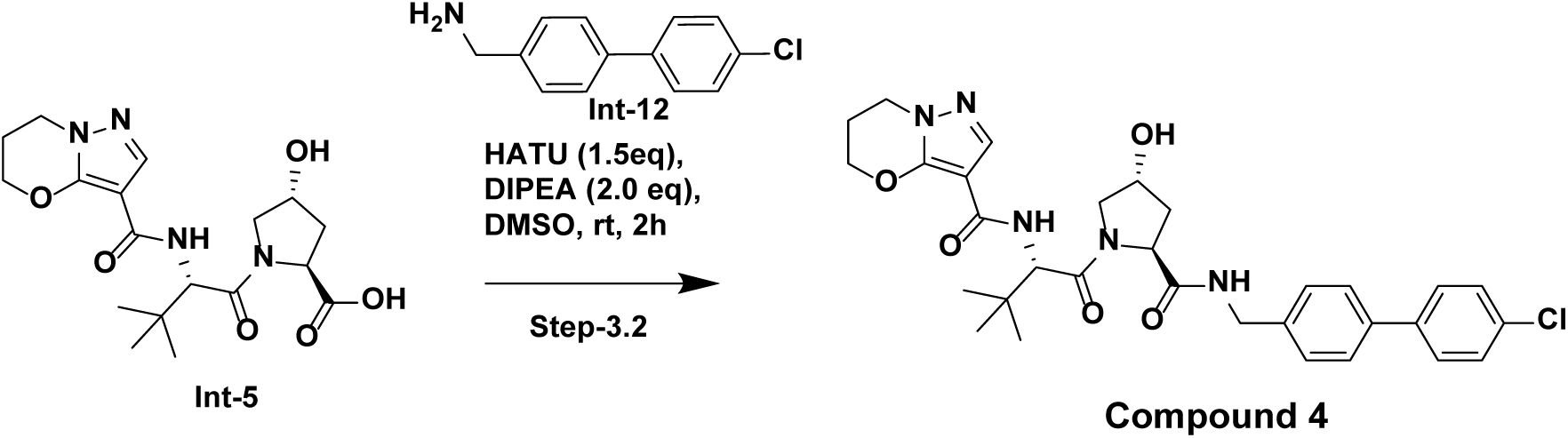

To a 4mL dram vial was added (2S,4R)-1-((S)-2-(6,7-dihydro-5H-pyrazolo[5,1-b][1,3]oxazine-3-carboxamido)-3,3-dimethylbutanoyl)-4-hydroxypyrrolidine-2-carboxylic acid (16 mg, 0.041 mmol), (4’-chloro-[1,1’-biphenyl]-4-yl)methanamine (8.83 mg, 0.041 mmol), N-ethyl-N-isopropylpropan-2-amine (7.86 mg, 10.63 µL, 0.061 mmol) 2-(3H-[1,2,3]triazolo[4,5-b]pyridin-3-yl)-1,1,3,3-tetramethylisouronium hexafluorophosphate(V) (30.8 mg, 0.081 mmol) in dimethyl sulfoxide (0.5 mL) and stirred at room temperature for 2 hours. The reaction mixture was purified by prep-HPLC on a Phenomenex Luna column, 5 micron, C8(2), 100 Å, 150 × 21.2 mm eluting at 20 mL/min with a gradient of 10-100% acetonitrile (0.1% TFA) in acetonitrile (0.1% TFA). The combined fractions containing the product were frozen at -78 °C, and lyophilized. N-((S)-1-((2S,4R)-2-(((4’-chloro-[1,1’-biphenyl]-4-yl)methyl)carbamoyl)-4-hydroxypyrrolidin-1-yl)-3,3-dimethyl-1-oxobutan-2-yl)-6,7-dihydro-5H-pyrazolo[5,1-b][1,3]oxazine-3-carboxamide (8.9 mg, 0.015 mmol, Yield: 36.9 %, Purity: 99%) obtained as a white solid.

LCMS performed on Agilent 1260 Infinity. 2 ml/min Gradient 5-95B (1.5 min); 95-95B (0.3 min); 95-5B (0.2 min) Injection Volume 1 uL Mobile Phase A H20 (0.1% TFA in Water) Mobile Phase B ACN (0.1% TFA in ACN) m/z: 594.248 (M+1), RT: 1.174min

^1^H NMR (400 MHz, DMSO-*d*_6_) δ ppm 8.52 (br t, *J*=5.9 Hz, 1 H), 7.64 - 7.72 (m, 2 H), 7.57 - 7.62 (m, 3 H), 7.49 - 7.54 (m, 2 H), 7.39 (d, *J*=8.4 Hz, 2 H), 6.82 (d, *J*=9.4 Hz, 1 H), 4.69 (br d, *J*=9.4 Hz, 1 H), 4.60 (br d, *J*=9.0 Hz, 1 H), 4.41 - 4.55 (m, 4 H), 4.26 - 4.39 (m, 5 H), 4.11 (br t, *J*=6.1 Hz, 4 H), 3.66 - 3.69 (m, 1 H), 2.55 (br s, 1 H), 2.24 (br d, *J*=3.3 Hz, 2 H), 2.00 - 2.11 (m, 1 H), 1.91 (ddd, *J*=12.9, 8.5, 4.5 Hz, 1 H), 0.93 - 1.02 (m, 9 H)

Step-3-2: Synthesis of N-((S)-1-((2S,4R)-2-(((2’-chloro-[1,1’-biphenyl]-4-yl)methyl)carbamoyl)-4-hydroxypyrrolidin-1-yl)-3,3-dimethyl-1-oxobutan--yl)-6,7-dihydro-5H-pyrazolo[5,1-b][1,3]oxazine-3-carboxamide (Compound 5):

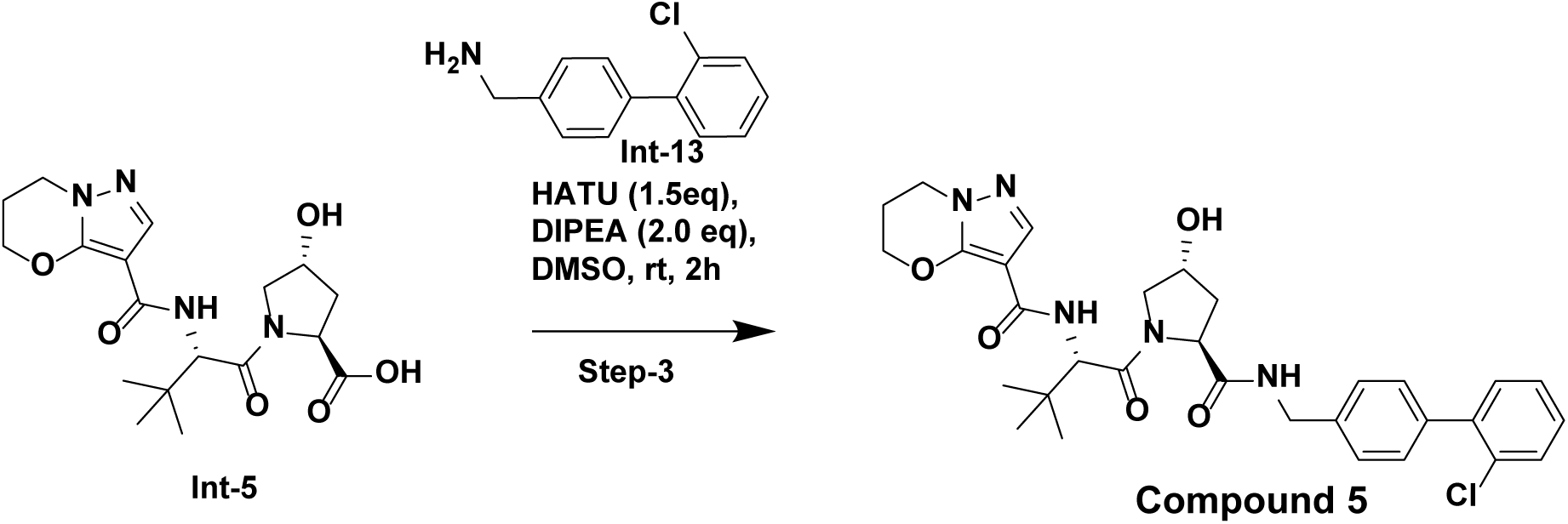

To a 4mL dram vial was added (2S,4R)-1-((S)-2-(6,7-dihydro-5H-pyrazolo[5,1-b][1,3]oxazine-3-carboxamido)-3,3-dimethylbutanoyl)-4-hydroxypyrrolidine-2-carboxylic acid (16 mg, 0.041 mmol), (2’-chloro-[1,1’-biphenyl]-4-yl)methanamine (8.83 mg, 0.041 mmol), 1,1’-dimethyltriethylamine (10.49 mg, 0.014 mL, 0.081 mmol), HATU (23.14 mg, 0.061 mmol) in dimethyl sulfoxide (.75mL) stirred at room temperature for 2 hours. The reaction mixture was purified by prep-HPLC on a Phenomenex Luna column, 5 micron, C8(2), 100 Å, 150 × 21.2 mm eluting at 20 mL/min with a gradient of 10-100% acetonitrile (0.1% TFA) in acetonitrile (0.1% TFA). The combined fractions containing the product were frozen at -78 °C, and lyophilized. N-((S)-1-((2S,4R)-2-(((2’-chloro-[1,1’-biphenyl]-4-yl)methyl)carbamoyl)-4-hydroxypyrrolidin-1-yl)-3,3-dimethyl-1-oxobutan-2-yl)-6,7-dihydro-5H-pyrazolo[5,1-b][1,3]oxazine-3-carboxamide (8.5 mg, 0.014 mmol, Yield: 35.3 %, Purity: 99%) obtained as a white solid.

LCMS performed on Agilent 1260 Infinity. 2 ml/min Gradient 5-95B (1.5 min); 95-95B (0.3 min); 95-5B (0.2 min) Injection Volume 1 uL Mobile Phase A H20 (0.1% TFA in Water) Mobile Phase B ACN (0.1% TFA in ACN) m/z: 594.248 (M+1), RT: 1.133min

^1^H NMR (400 MHz, DMSO-*d*_6_) δ ppm 8.55 (br t, *J*=6.0 Hz, 1 H), 7.60 (s, 1 H), 7.53 - 7.60 (m, 1 H), 7.34 - 7.46 (m, 7 H), 6.82 (d, *J*=9.2 Hz, 1 H), 4.69 (d, *J*=9.4 Hz, 1 H), 4.61 (br d, *J*=9.0 Hz, 1 H), 4.29 - 4.54 (m, 8 H), 4.00 - 4.20 (m, 5 H), 3.70 - 3.90 (m, 1 H), 3.66 - 3.68 (m, 1 H), 2.55 (br s, 1 H), 2.21 - 2.27 (m, 2 H), 2.02 - 2.11 (m, 1 H), 1.93 (ddd, *J*=12.9, 8.6, 4.5 Hz, 1 H), 0.93 - 1.01 (m, 9 H)

Step-3-3: Synthesis N-((S)-1-((2S,4R)-2-(([1,1’-biphenyl]-4-ylmethyl)carbamoyl)-4-hydroxypyrrolidin-1-yl)-3,3- dimethyl-1-oxobutan-2-yl)-6,7-dihydro-5H-pyrazolo[5,1-b][1,3]oxazine-3-carboxamide (Compound 6)

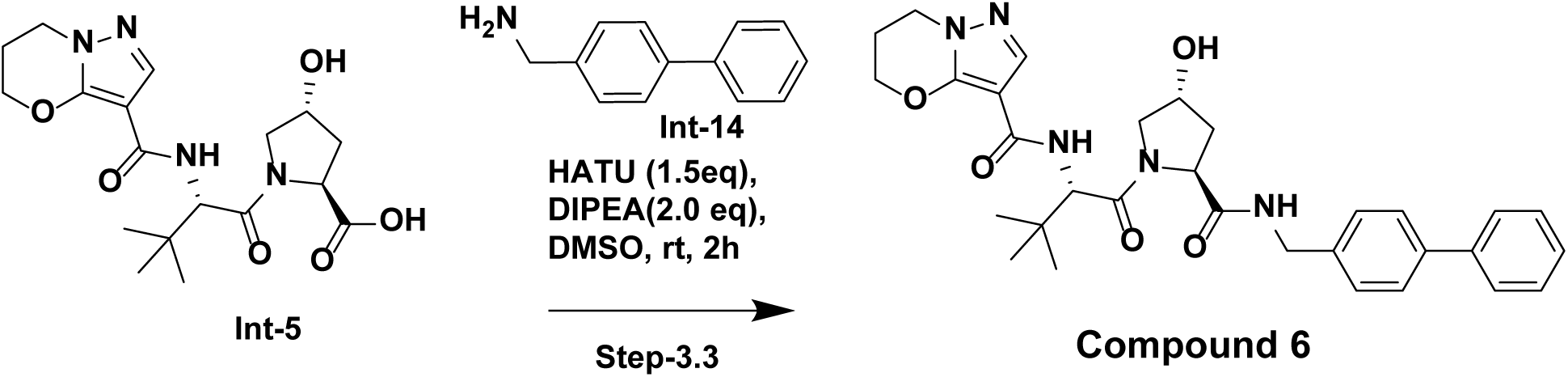

To a 4mL dram vial was added tert-butyl (2S,4R)-1-((S)-2-(6,7-dihydro-5H-pyrazolo[5,1-b][1,3]oxazine-3-carboxamido)-3,3-dimethylbutanoyl)-4-hydroxypyrrolidine-2-carboxylate (20 mg, 0.044 mmol, 134023-45), [1,1’-biphenyl]-4-ylmethanamine (9.76 mg, 0.053 mmol), N-ethyl-N-isopropylpropan-2-amine (11.47 mg, 0.016 mL, 0.089 mmol) in dimethyl sulfoxide (0.75 mL) and stirred at room temperature for 2 hours. The reaction mixture was purified by prep-HPLC on a Phenomenex Luna column, 5 micron, C8(2), 100 Å, 150 × 21.2 mm eluting at 20 mL/min with a gradient of 10-100% acetonitrile (0.1% TFA) in acetonitrile (0.1% TFA). The combined fractions containing the product were frozen at -78 °C, and lyophilized. N-((S)-1-((2S,4R)-2-(([1,1’-biphenyl]-4-ylmethyl)carbamoyl)-4-hydroxypyrrolidin-1-yl)-3,3-dimethyl-1-oxobutan-2-yl)-6,7-dihydro-5H-pyrazolo[5,1-b][1,3]oxazine-3-carboxamide (16.0 mg, 0.029 mmol, Yield: 64.4 %, Purity: 99%) obtained as a white solid.

LCMS performed on Agilent 1260 Infinity. 2 ml/min Gradient 5-95B (1.5 min); 95-95B (0.3 min); 95-5B (0.2 min) Injection Volume 1 uL Mobile Phase A H20 (0.1% TFA in Water) Mobile Phase B ACN (0.1% TFA in ACN) m/z: 560.287 (M+1), RT: 1.089min

^1^H NMR (400 MHz, DMSO-*d*_6_) δ ppm 7.65 (d, *J*=7.3 Hz, 2 H), 7.58 - 7.62 (m, 3 H), 7.47 (t, *J*=7.6 Hz, 2 H), 7.34 - 7.41 (m, 3 H), 4.52 (br s, 2 H), 4.30 - 4.37 (m, 3 H), 4.10 (t, *J*=6.1 Hz, 2 H), 3.69 - 3.80 (m, 2 H), 3.66 - 3.68 (m, 1 H), 2.24 (br d, *J*=2.9 Hz, 2 H), 0.95 - 1.00 (m, 9 H)

## Author Contributions

Conceptualization: J.W.B., S.-C.O., T.D., E.I., L.L., J.M.N., M.D.H., A.C., J.I.T., A.Ve., A.B., G.S.P., P.A.T., S.B., O.B., J.M., and P.R.P.; Methodology: J.W.B., W.D., K.T.G.S., H.-Y.L., S.L., A.Va., A.G., S.-C.O., J.A., R.K.H., H.H., T.D., E.I., L.L., J.M.N., M.D.H., A.R., C.-C.L., A.C., and J.I.T.; Investigation: J.W.B., W.D., K.T.G.S., H.-Y.L., S.L., A.Va., T.D., E.I., J.W., R.R., L.L., J.M.N., M.D.H., A.R., A.D.S., C.-C.L., D.T., G.B., and J.I.T.; Formal Analysis: J.W.B., W.D., K.T.G.S., H.-Y.L., S.L., A.Va., S.-C.O., T.D., E.I., J.W., R.R., L.L., J.M.N., M.D.H., A.R., A.D.S., C.-C.L., D.T., A.C., G.B., and J.I.T.; Data Curation: J.W.B., W.D., J.W., R.R., A.R., A.D.S., C.-C.L., D.T., A.C., and G.B.; Resources: J.W.B., A.G., J.A., R.K.H., H.H., W.d.B., S.V., B.Z., and K.S.A.; Visualization: J.W.B.; Supervision: A.Ve., A.B., G.S.P., P.A.T., S.B., W.d.B., S.V., B.Z., K.S.A., O.B., J.M., and P.R.P.; Writing – original draft: J.W.B.; Writing – review & editing: All authors.

## Data Availability Statement

Data supporting the findings in this study are provided in the main text and supplementary materials. The compound screen identifying dGEM3 was performed on Plexium’s proprietary screening platform. Mass spectrometry raw data files have been deposited to the PRIDE Archive (PDX######). Tables corresponding to RNA-seq, TurboID, BioE3, and global proteomics experiments are available in Supplementary Data Set 1.

## Competing interests

Financial support for this research was provided by Amgen and Plexium. All authors were employed by Amgen or Plexium while this study was conducted, as indicated in the affiliations.

## Supporting information

Supplementary Data Set 1

## Acknowledgements

We sincerely thank the teams at Plexium and Amgen for their invaluable collaboration and contributions to this research. We thank the technology team at Plexium for the development of the Picowell RNAseq screening platform. We also thank Kyle Mangano, Emily Cook, Sung Jun Bae, Kate Smither, Jianzhong Hu, Amine Sadok and all members of the Induced Proximity Platform and Amgen R&D Postdoctoral Fellows Program for thoughtful scientific discussions and feedback on this work. Support from the Research Materials Management and Structural Biology teams is also greatly appreciated. Schematics were created with the use of BioRender.com.

## References

1 Sakamoto, K. M. et al. Protacs: chimeric molecules that target proteins to the Skp1-Cullin-F box complex for ubiquitination and degradation. Proc Natl Acad Sci U S A 98, 8554–8559 (2001). 10.1073/pnas.141230798

2 Schneekloth, A. R., Pucheault, M., Tae, H. S. & Crews, C. M. Targeted intracellular protein degradation induced by a small molecule: En route to chemical proteomics. Bioorganic & Medicinal Chemistry Letters 18, 5904–5908 (2008). 10.1016/j.bmcl.2008.07.114

3 Winter, G. E. et al. Phthalimide conjugation as a strategy for in vivo target protein degradation. Science 348, 1376–1381 (2015). 10.1126/science.aab1433

4 Zengerle, M., Chan, K.-H. & Ciulli, A. Selective Small Molecule Induced Degradation of the BET Bromodomain Protein BRD4. ACS Chemical Biology 10, 1770–1777 (2015). 10.1021/acschembio.5b00216

5 Diehl, C. J. & Ciulli, A. Discovery of small molecule ligands for the von Hippel-Lindau (VHL) E3 ligase and their use as inhibitors and PROTAC degraders. Chem Soc Rev 51, 8216–8257 (2022). 10.1039/d2cs00387b

6 Tsai, J. M., Nowak, R. P., Ebert, B. L. & Fischer, E. S. Targeted protein degradation: from mechanisms to clinic. Nat Rev Mol Cell Biol 25, 740–757 (2024). 10.1038/s41580-024-00729-9

7 Bekes, M., Langley, D. R. & Crews, C. M. PROTAC targeted protein degraders: the past is prologue. Nat Rev Drug Discov 21, 181–200 (2022). 10.1038/s41573-021-00371-6

8 Starodub, A. et al. Phase 1 study of KT-333, a targeted protein degrader, in patients with relapsed or refractory lymphomas, large granular lymphocytic leukemia, and solid tumors. Journal of Clinical Oncology 40, TPS3171-TPS3171 (2022). 10.1200/JCO.2022.40.16_suppl.TPS3171

9 Sharma, K. et al. Abstract LB037: E3 pairing and structural mechanisms underlying anti-tumor activity of clinical STAT3 degrader KT-333. Cancer Research 84, LB037-LB037 (2024). 10.1158/1538-7445.Am2024-lb037

10 Thummuri, D. et al. Overcoming Gemcitabine Resistance in Pancreatic Cancer Using the BCL-XL–Specific Degrader DT2216. Molecular Cancer Therapeutics 21, 184–192 (2022). 10.1158/1535-7163.Mct-21-0474

11 Domostegui, A., Nieto-Barrado, L., Perez-Lopez, C. & Mayor-Ruiz, C. Chasing molecular glue degraders: screening approaches. Chem Soc Rev 51, 5498–5517 (2022). 10.1039/d2cs00197g

12 Kannt, A. & Dikic, I. Expanding the arsenal of E3 ubiquitin ligases for proximity-induced protein degradation. Cell Chem Biol 28, 1014-1031 (2021). 10.1016/j.chembiol.2021.04.007

13 Ito, T. et al. Identification of a primary target of thalidomide teratogenicity. Science 327, 1345–1350 (2010). 10.1126/science.1177319

14 Fischer, E. S. et al. Structure of the DDB1–CRBN E3 ubiquitin ligase in complex with thalidomide. Nature 512, 49–53 (2014). 10.1038/nature13527

15 Chamberlain, P. P. et al. Structure of the human Cereblon–DDB1–lenalidomide complex reveals basis for responsiveness to thalidomide analogs. Nature Structural & Molecular Biology 21, 803–809 (2014). 10.1038/nsmb.2874

16 Kronke, J. et al. Lenalidomide causes selective degradation of IKZF1 and IKZF3 in multiple myeloma cells. Science 343, 301–305 (2014). 10.1126/science.1244851

17 Lu, G. et al. The myeloma drug lenalidomide promotes the cereblon-dependent destruction of Ikaros proteins. Science 343, 305–309 (2014). 10.1126/science.1244917

18 Donovan, K. A. et al. Thalidomide promotes degradation of SALL4, a transcription factor implicated in Duane Radial Ray syndrome. eLife 7 (2018). 10.7554/eLife.38430

19 Matyskiela, M. E. et al. Crystal structure of the SALL4-pomalidomide-cereblon-DDB1 complex. Nat Struct Mol Biol 27, 319–322 (2020). 10.1038/s41594-020-0405-9

20 Han, T. et al. Anticancer sulfonamides target splicing by inducing RBM39 degradation via recruitment to DCAF15. Science 356 (2017). 10.1126/science.aal3755

21 Uehara, T. et al. Selective degradation of splicing factor CAPERalpha by anticancer sulfonamides. Nat Chem Biol 13, 675–680 (2017). 10.1038/nchembio.2363

22 Faust, T. B. et al. Structural complementarity facilitates E7820-mediated degradation of RBM39 by DCAF15. Nature Chemical Biology 16, 7–14 (2019). 10.1038/s41589-019-0378-3

23 Bussiere, D. E. et al. Structural basis of indisulam-mediated RBM39 recruitment to DCAF15 E3 ligase complex. Nature Chemical Biology 16, 15–23 (2019). 10.1038/s41589-019-0411-6

24 Du, X. et al. Structural Basis and Kinetic Pathway of RBM39 Recruitment to DCAF15 by a Sulfonamide Molecular Glue E7820. Structure 27, 1625–1633 e1623 (2019). 10.1016/j.str.2019.10.005

25 Bettayeb, K. et al. CR8, a potent and selective, roscovitine-derived inhibitor of cyclin-dependent kinases. Oncogene 27, 5797–5807 (2008). 10.1038/onc.2008.191

26 Slabicki, M. et al. The CDK inhibitor CR8 acts as a molecular glue degrader that depletes cyclin K. Nature 585, 293–297 (2020). 10.1038/s41586-020-2374-x

27 Mayor-Ruiz, C. et al. Rational discovery of molecular glue degraders via scalable chemical profiling. Nat Chem Biol 16, 1199–1207 (2020). 10.1038/s41589-020-0594-x

28 Lv, L. et al. Discovery of a molecular glue promoting CDK12-DDB1 interaction to trigger cyclin K degradation. Elife 9 (2020). 10.7554/eLife.59994

29 Kozicka, Z. et al. Design principles for cyclin K molecular glue degraders. Nat Chem Biol 20, 93–102 (2024). 10.1038/s41589-023-01409-z

30 Tutter, A. et al. A small molecule VHL molecular glue degrader for cysteine dioxygenase 1. bioRxiv (2024). 10.1101/2024.01.25.576086

31 Frost, J. et al. Potent and selective chemical probe of hypoxic signalling downstream of HIF-alpha hydroxylation via VHL inhibition. Nat Commun 7, 13312 (2016). 10.1038/ncomms13312

32 Filippakopoulos, P. et al. Selective inhibition of BET bromodomains. Nature 468, 1067–1073 (2010). 10.1038/nature09504

33 Kerres, N. et al. Chemically Induced Degradation of the Oncogenic Transcription Factor BCL6. Cell Rep 20, 2860–2875 (2017). 10.1016/j.celrep.2017.08.081

34 Slabicki, M. et al. Small-molecule-induced polymerization triggers degradation of BCL6. Nature 588, 164–168 (2020). 10.1038/s41586-020-2925-1

35 Shergalis, A. G. et al. CRISPR Screen Reveals BRD2/4 Molecular Glue-like Degrader via Recruitment of DCAF16. ACS Chem Biol 18, 331–339 (2023). 10.1021/acschembio.2c00747

36 Sarott, R. C. et al. Chemical Specification of E3 Ubiquitin Ligase Engagement by Cysteine-Reactive Chemistry. J Am Chem Soc 145, 21937–21944 (2023). 10.1021/jacs.3c06622

37 Li, Y. D. et al. Template-assisted covalent modification underlies activity of covalent molecular glues. Nat Chem Biol (2024). 10.1038/s41589-024-01668-4

38 Hassan, M. M. et al. Exploration of the Tunability of BRD4 Degradation by DCAF16 Trans-labelling Covalent Glues. BioRxiv (2023). 10.1101/2023.10.07.561308

39 Zhuang, Z. et al. Discovery of electrophilic degraders that exploit SNAr chemistry. bioRxiv, 2024.2009.2025.615094 (2024). 10.1101/2024.09.25.615094

40 Toriki, E. S. et al. Rational Chemical Design of Molecular Glue Degraders. ACS Cent Sci 9, 915–926 (2023). 10.1021/acscentsci.2c01317

41 Parker, G. S. et al. Discovery of Monovalent Direct Degraders of BRD4 that Act via the Recruitment of DCAF11. Molecular Cancer Therapeutics 23, 1446–1458 (2024). 10.1158/1535-7163.Mct-24-0219

42 Oleinikovas, V., Gainza, P., Ryckmans, T., Fasching, B. & Thoma, N. H. From Thalidomide to Rational Molecular Glue Design for Targeted Protein Degradation. Annu Rev Pharmacol Toxicol (2023). 10.1146/annurev-pharmtox-022123-104147

43 Matyskiela, M. E. et al. A novel cereblon modulator recruits GSPT1 to the CRL4(CRBN) ubiquitin ligase. Nature 535, 252–257 (2016). 10.1038/nature18611

44 Wang, E. S. et al. Acute pharmacological degradation of Helios destabilizes regulatory T cells. Nat Chem Biol 17, 711–717 (2021). 10.1038/s41589-021-00802-w

45 Bonazzi, S. et al. Discovery and characterization of a selective IKZF2 glue degrader for cancer immunotherapy. Cell Chem Biol 30, 235–247 e212 (2023). 10.1016/j.chembiol.2023.02.005

46 Miyamoto, D. K. et al. Design and Development of IKZF2 and CK1alpha Dual Degraders. J Med Chem 66, 16953–16979 (2023). 10.1021/acs.jmedchem.3c01736

47 Ng, A. et al. Discovery of Molecular Glue Degraders via Isogenic Morphological Profiling. ACS Chem Biol 18, 2464–2473 (2023). 10.1021/acschembio.3c00598

48 Gadd, M. S. et al. Structural basis of PROTAC cooperative recognition for selective protein degradation. Nature Chemical Biology 13, 514–521 (2017). 10.1038/nchembio.2329

49 Nowak, R. P. et al. Plasticity in binding confers selectivity in ligand-induced protein degradation. Nature Chemical Biology 14, 706–714 (2018). 10.1038/s41589-018-0055-y

50 Branon, T. C. et al. Efficient proximity labeling in living cells and organisms with TurboID. Nat Biotechnol 36, 880–887 (2018). 10.1038/nbt.4201

51 Yamanaka, S. et al. A proximity biotinylation-based approach to identify protein-E3 ligase interactions induced by PROTACs and molecular glues. Nat Commun 13, 183 (2022). 10.1038/s41467-021-27818-z

52 Tsuiji, H. et al. Spliceosome integrity is defective in the motor neuron diseases ALS and SMA. EMBO Molecular Medicine 5, 221–234 (2013). 10.1002/emmm.201202303

53 Barroso-Gomila, O. et al. BioE3 identifies specific substrates of ubiquitin E3 ligases. Nat Commun 14, 7656 (2023). 10.1038/s41467-023-43326-8

54 Huang, H. T. et al. Ubiquitin-specific proximity labeling for the identification of E3 ligase substrates. Nat Chem Biol (2024). 10.1038/s41589-024-01590-9

55 Frost, J., Rocha, S. & Ciulli, A. Von Hippel-Lindau (VHL) small-molecule inhibitor binding increases stability and intracellular levels of VHL protein. J Biol Chem 297, 100910 (2021). 10.1016/j.jbc.2021.100910

56 Curmi, F. & Cauchi, R. J. The multiple lives of DEAD-box RNA helicase DP103/DDX20/Gemin3. Biochem Soc Trans 46, 329–341 (2018). 10.1042/BST20180016

57 Panek, J. et al. The SMN complex drives structural changes in human snRNAs to enable snRNP assembly. Nat Commun 14, 6580 (2023). 10.1038/s41467-023-42324-0

58 Jumper, J. et al. Highly accurate protein structure prediction with AlphaFold. Nature 596, 583–589 (2021). 10.1038/s41586-021-03819-2

59 Charroux, B. et al. Gemin3: A Novel Dead Box Protein That Interacts with Smn, the Spinal Muscular Atrophy Gene Product, and Is a Component of Gems. Journal of Cell Biology 147, 1181–1194 (1999). 10.1083/jcb.147.6.1181

60 Klappacher, G. W. et al. An Induced Ets Repressor Complex Regulates Growth Arrest during Terminal Macrophage Differentiation. Cell 109, 169–180 (2002). 10.1016/S0092-8674(02)00714-6

61 Yan, X., Mouillet, J. F., Ou, Q. & Sadovsky, Y. A novel domain within the DEAD-box protein DP103 is essential for transcriptional repression and helicase activity. Mol Cell Biol 23, 414–423 (2003). 10.1128/MCB.23.1.414-423.2003

62 van Kempen, M. et al. Fast and accurate protein structure search with Foldseek. Nat Biotechnol (2023). 10.1038/s41587-023-01773-0

63 Schütz, P. et al. Comparative Structural Analysis of Human DEAD-Box RNA Helicases. PLOS ONE 5, e12791 (2010). 10.1371/journal.pone.0012791

64 Dove, K. K. et al. Two functionally distinct E2/E3 pairs coordinate sequential ubiquitination of a common substrate in Caenorhabditis elegans development. Proc Natl Acad Sci U S A 114, E6576–E6584 (2017). 10.1073/pnas.1705060114

65 Scott, D. C. et al. Two Distinct Types of E3 Ligases Work in Unison to Regulate Substrate Ubiquitylation. Cell 166, 1198–1214.e1124 (2016). 10.1016/j.cell.2016.07.027

66 Roy, M. J. et al. SPR-Measured Dissociation Kinetics of PROTAC Ternary Complexes Influence Target Degradation Rate. ACS Chem Biol 14, 361–368 (2019). 10.1021/acschembio.9b00092

67 Mason, J. W. et al. DNA-encoded library-enabled discovery of proximity-inducing small molecules. Nat Chem Biol (2023). 10.1038/s41589-023-01458-4

68 Xiong, Y. et al. Chemo-proteomics exploration of HDAC degradability by small molecule degraders. Cell Chem Biol 28, 1514–1527 e1514 (2021). 10.1016/j.chembiol.2021.07.002

69 Baek, K. et al. Unveiling the hidden interactome of CRBN molecular glues with chemoproteomics. bioRxiv (2024). 10.1101/2024.09.11.612438

70 Fan, A. T. et al. A Kinetic Scout Approach Accelerates Targeted Protein Degrader Development. bioRxiv, 2024.2009.2017.612508 (2024). 10.1101/2024.09.17.612508

71 Mayor-Ruiz, C. et al. Plasticity of the Cullin-RING Ligase Repertoire Shapes Sensitivity to Ligand-Induced Protein Degradation. Mol Cell 75, 849–858 e848 (2019). 10.1016/j.molcel.2019.07.013

72 Hill, S. et al. Robust cullin-RING ligase function is established by a multiplicity of poly-ubiquitylation pathways. Elife 8 (2019). 10.7554/eLife.51163

73 Reitsma, J. M. et al. Composition and Regulation of the Cellular Repertoire of SCF Ubiquitin Ligases. Cell 171, 1326–1339 e1314 (2017). 10.1016/j.cell.2017.10.016

74 Reichermeier, K. M. et al. PIKES Analysis Reveals Response to Degraders and Key Regulatory Mechanisms of the CRL4 Network. Mol Cell 77, 1092–1106 e1099 (2020). 10.1016/j.molcel.2019.12.013

75 Rosenberg, S. C. et al. Ternary complex dissociation kinetics contribute to mutant-selective EGFR degradation. Cell Chem Biol (2023). 10.1016/j.chembiol.2023.01.007

76 Wurz, R. P. et al. Affinity and cooperativity modulate ternary complex formation to drive targeted protein degradation. Nat Commun 14, 4177 (2023). 10.1038/s41467-023-39904-5

77 R: A Language and Environment for Statistical Computing (2023).

78 Martin, M. Cutadapt removes adapter sequences from high-throughput sequencing reads. 2011 17, 3 (2011). 10.14806/ej.17.1.200

79 Dobin, A. et al. STAR: ultrafast universal RNA-seq aligner. Bioinformatics 29, 15–21 (2012). 10.1093/bioinformatics/bts635

80 Liao, Y., Smyth, G. K. & Shi, W. featureCounts: an efficient general purpose program for assigning sequence reads to genomic features. Bioinformatics 30, 923–930 (2013). 10.1093/bioinformatics/btt656

81 Love, M. I., Huber, W. & Anders, S. Moderated estimation of fold change and dispersion for RNA-seq data with DESeq2. Genome Biology 15, 550 (2014). 10.1186/s13059-014-0550-8

82 da Veiga Leprevost, F., et al. Philosopher: a versatile toolkit for shotgun proteomics data analysis. Nat Methods 17, 869–870 (2020). 10.1038/s41592-020-0912-y

83 Yu, F., Haynes, S. E. & Nesvizhskii, A. I. IonQuant Enables Accurate and Sensitive Label-Free Quantification With FDR-Controlled Match-Between-Runs. Mol Cell Proteomics 20, 100077 (2021). 10.1016/j.mcpro.2021.100077

84 Ritchie, M. E. et al. limma powers differential expression analyses for RNA-sequencing and microarray studies. Nucleic Acids Research 43, e47–e47 (2015). 10.1093/nar/gkv007

85 Békés, M. et al. DUB-Resistant Ubiquitin to Survey Ubiquitination Switches in Mammalian Cells. Cell Reports 5, 826–838 (2013). 10.1016/j.celrep.2013.10.008

86 Demichev, V., Messner, C. B., Vernardis, S. I., Lilley, K. S. & Ralser, M. DIA-NN: neural networks and interference correction enable deep proteome coverage in high throughput. Nature Methods 17, 41–44 (2020). 10.1038/s41592-019-0638-x

87 Demichev, V. et al. dia-PASEF data analysis using FragPipe and DIA-NN for deep proteomics of low sample amounts. Nat Commun 13, 3944 (2022). 10.1038/s41467-022-31492-0

88 Skowronek, P. et al. Rapid and In-Depth Coverage of the (Phospho-)Proteome With Deep Libraries and Optimal Window Design for dia-PASEF. Molecular & Cellular Proteomics 21 (2022). 10.1016/j.mcpro.2022.100279

89 Choi, M. et al. MSstats: an R package for statistical analysis of quantitative mass spectrometry-based proteomic experiments. Bioinformatics 30, 2524–2526 (2014). 10.1093/bioinformatics/btu305

90 Canny, M. D. et al. Inhibition of 53BP1 favors homology-dependent DNA repair and increases CRISPR– Cas9 genome-editing efficiency. Nature Biotechnology 36, 95–102 (2018). 10.1038/nbt.4021

